# Supramolecular Organization Predicts Protein Nanoparticle Delivery to Neutrophils for Acute Lung Inflammation Diagnosis and Treatment

**DOI:** 10.1101/2020.04.15.037564

**Authors:** Jacob W. Myerson, Priyal N. Patel, Nahal Habibi, Landis R. Walsh, Yi-Wei Lee, David C. Luther, Laura T. Ferguson, Michael H. Zaleski, Marco E. Zamora, Oscar A. Marcos-Contreras, Patrick M. Glassman, Ian Johnston, Elizabeth D. Hood, Tea Shuvaeva, Jason V. Gregory, Raisa Y. Kiseleva, Jia Nong, Kathryn M. Rubey, Colin F. Greineder, Samir Mitragotri, George S. Worthen, Vincent M. Rotello, Joerg Lahann, Vladimir R. Muzykantov, Jacob S. Brenner

## Abstract

Acute lung inflammation has severe morbidity, as seen in COVID-19 patients. Lung inflammation is accompanied or led by massive accumulation of neutrophils in pulmonary capillaries (“margination”). We sought to identify nanostructural properties that predispose nanoparticles to accumulate in pulmonary marginated neutrophils, and therefore to target severely inflamed lungs. We designed a library of nanoparticles and conducted an *in vivo* screen of biodistributions in naive mice and mice treated with lipopolysaccharides. We found that supramolecular organization of protein in nanoparticles predicts uptake in inflamed lungs. Specifically, nanoparticles with agglutinated protein (NAPs) efficiently home to pulmonary neutrophils, while protein nanoparticles with symmetric structure (*e.g.* viral capsids) are ignored by pulmonary neutrophils. We validated this finding by engineering protein-conjugated liposomes that recapitulate NAP targeting to neutrophils in inflamed lungs. We show that NAPs can diagnose acute lung injury in SPECT imaging and that NAP-like liposomes can mitigate neutrophil extravasation and pulmonary edema arising in lung inflammation. Finally, we demonstrate that ischemic *ex vivo* human lungs selectively take up NAPs, illustrating translational potential. This work demonstrates that structure-dependent interactions with neutrophils can dramatically alter the biodistribution of nanoparticles, and NAPs have significant potential in detecting and treating respiratory conditions arising from injury or infections.

---

The COVID-19 pandemic tragically illustrates the dangers of acute inflammation and infection of the lungs, for both individuals and societies. Acute alveolar inflammation causes the clinical syndrome known as acute respiratory distress syndrome (ARDS), in which inflammation prevents the lungs from oxygenating the blood. Severe ARDS is the cause of death in most COVID-19 mortality and was a major cause of death in the 1918 influenza epidemic, but ARDS is common even outside of epidemics, affecting ∼200,000 American patients per year with a ∼35-50% mortality rate.^1-6^ ARDS is caused not just by viral infections, but also by sepsis, pneumonia (viral and bacterial), aspiration, and trauma.^3,4^ Largely because ARDS patients have poor tolerance of drug side effects, no pharmacological strategy has succeeded as an ARDS treatment.^5,7-9^ Therefore, there is an urgent need to develop drug delivery strategies that specifically target inflamed alveoli in ARDS and minimize systemic side effects.

Neutrophils are “first responder” cells in acute inflammation, rapidly adhering and activating in large numbers in inflamed vessels and forming populations of “marginated” neutrophils along the vascular lumen.^10-16^ Neutrophils can be activated by a variety of initiating factors, including pathogen- and damage-associated molecular patterns such as bacterial lipopolysaccharides (LPS).^17,18^ After acute inflammatory insults, neutrophils marginate in most organs, but by far most avidly in the lung capillaries.^6,14,15,19,20^ Neutrophils are therefore key cell types in most forms of ARDS. In ARDS, marginated neutrophils can secrete tissue-damaging substances (proteases, reactive oxygen species) and extravasate into the alveoli, leading to disruption of the endothelial barrier and accumulation of neutrophils and edematous fluid in the air space of the lungs (Figure 1A).^6,15,16,19-21^

**Figure 1.**
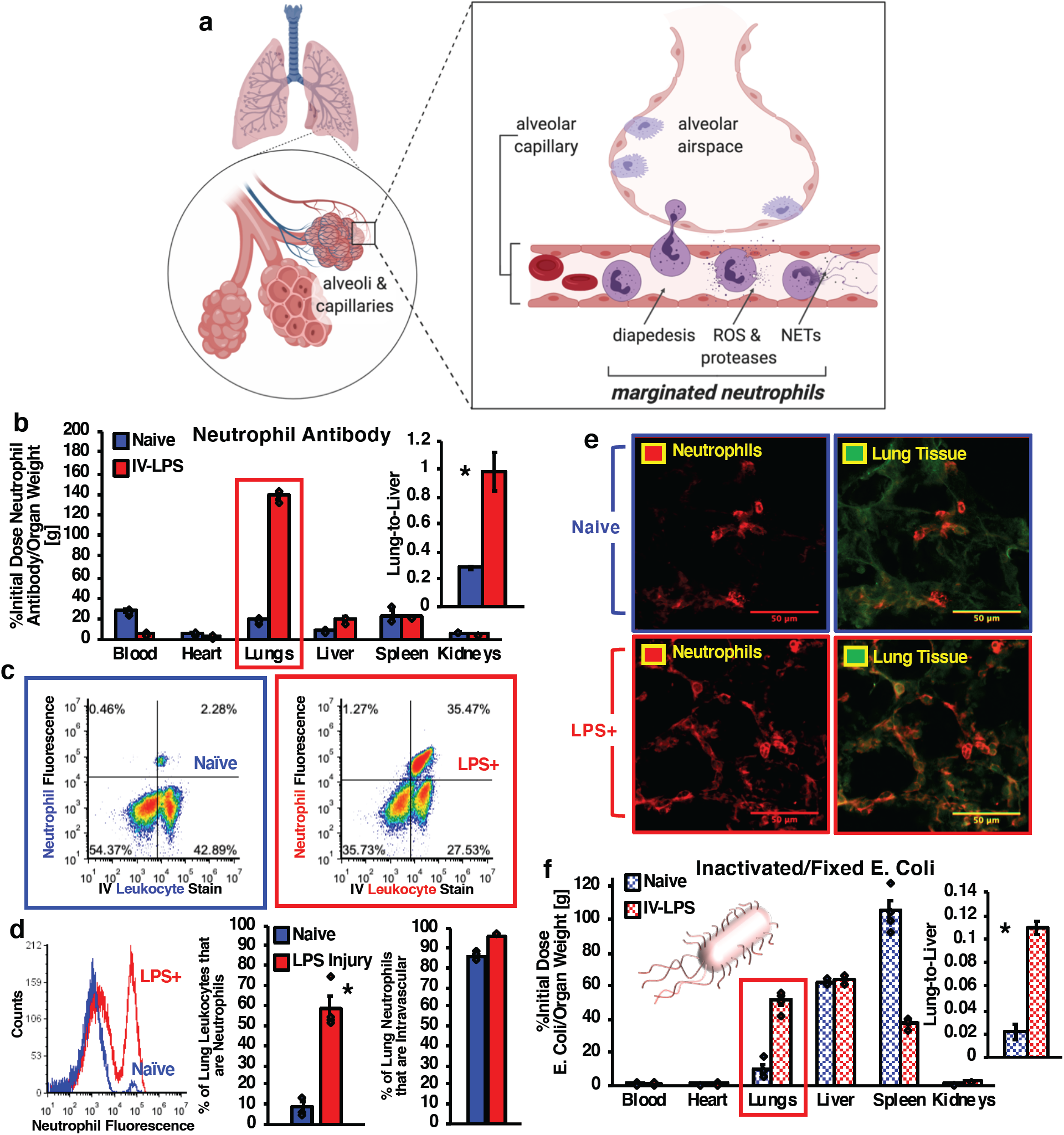
Neutrophil Accumulation in Acutely Inflamed Pulmonary Vasculature. (a) Schematic of neutrophil margination and extravasation in inflamed lungs. (b) Biodistributions of anti-Ly6G antibody in naïve (n=3) and IV-LPS-injured (n=3) mice (*red box = p*<.*001, * = p*<.*01*). (c-d) Flow cytometric characterization of single cell suspensions prepared from naïve and IV-LPS-injured injured mouse lungs. (c) Vertical axis indicates Ly6G staining for neutrophils (APC signal) and horizontal axis indicates stain induced by intravenous anti-CD45 antibody (FITC signal). (d) Flow cytometry data indicating increased neutrophil concentration in IV-LPS-injured mouse lungs (red), compared to naïve lungs (blue), and correlation of *intravenous* leukocyte staining with neutrophils. (n=3, *\* = p*<.*01*). (e) Fluorescence micrographs indicating increased concentration of neutrophils in IV-LPS-injured mouse lungs. Red: anti-Ly6G stain. Green: tissue autofluorescence. (f) Biodistributions of heat-inactivated, fixed, and ^125^I-labeled *E. coli* in naïve (n=4) and IV-LPS-injured (n=4) mice (*red box = p*<.*001, * = p*<.*01*).

Targeted nanoparticle delivery to marginated neutrophils could provide an ARDS treatment with minimal side effects, but specific delivery to marginated neutrophils remains an open challenge. Antibodies against markers such as Ly6G have achieved targeting to neutrophils in mice, but also deplete populations of circulating neutrophils.^22-25^ Additionally, while Ly6G readily marks neutrophils in mice, there is no analogous specific and ubiquitous marker on human neutrophils.^22^ Therefore, antibody targeting strategies have not been widely adopted for targeted drug delivery to these cells.^24^ As another route to neutrophil targeting, two previous studies noted that activated neutrophils take up denatured and agglutinated bovine albumin, concluding that *denatured* protein was critical in neutrophil-particle interactions.^26,27^

Nanoparticle structural properties such as shape, size, and deformability can define unique targeting behaviors.^28-32^ Here, we screened a diverse panel of nanoparticles to determine the nanostructural properties that predict uptake in pulmonary marginated neutrophils during acute inflammation. As a high-throughput animal model for ARDS, we administered LPS to mice, causing a massive increase in pulmonary marginated neutrophils. We show that two initial leads in our screen, lysozyme-dextran nanogels (LDNGs) and crosslinked albumin nanoparticles (ANPs), selectively home to marginated neutrophils in inflamed lungs, but not naïve lungs. In our subsequent screen of over 20 diverse nanoparticles, we find that 13 protein nanoparticles, all defined by *agglutination* of protein in *amorphous* nanostructures (nanoparticles with agglutinated proteins, NAPs), but *not* by denatured protein, have specificity for LPS-inflamed lungs. In contrast to NAPs, we demonstrate that three *symmetric* protein nanostructures (viruses/nanocages) have biodistributions unaffected by LPS injury. We show that polystyrene nanoparticles and five liposome formulations do not accumulate in injured lungs, indicating that nanostructures that are not based on protein are not intrinsically drawn to marginated neutrophils in acute inflammation. We then engineered liposomes (the most clinically relevant nanoparticle drug carriers) as NAPs, through conjugation to protein modified with hydrophobic cyclooctynes, encouraging protein agglutination on the liposome surface by hydrophobic interactions. We thus show that *supramolecular organization* of proteins, rather than chemical composition, best predicts uptake in marginated neutrophils in acutely inflamed lungs.

We then demonstrate proof of concept for NAPs as diagnostic and therapeutic tools for ARDS. We show; a) ^111^In-labeled NAPs provide diagnostic imaging contrast that distinguishes inflammatory lung injury from cardiogenic pulmonary edema; b) NAP-liposomes can significantly ameliorate edema in a mouse model of severe ARDS; c) NAPs, but not crystalline protein nanostructures, accumulate in *ex vivo* human lungs rejected for transplant due to ARDS-like conditions.

Collectively, our results will demonstrate that supramolecular organization of protein, namely protein agglutination, predicts strong, intrinsic nanoparticle tropism for marginated neutrophils. This finding indicates that NAPs, encompassing a wide range of nanoparticles based on or incorporating protein, have biodistributions that are responsive to inflammation. NAPs could be useful beyond ARDS, since marginated neutrophils play a pathogenic role in a diverse array of inflammatory diseases, including infections, heart attack, and stroke.^10-14^ But our findings provide a clear path forward for using NAPs to improve diagnosis and treatment of ARDS.

## Characterization of Marginated Neutrophil Accumulation in Inflamed Lung Vasculature

To quantify the increase in pulmonary marginated neutrophils after inflammatory lung injury, radiolabeled clone 1A8 anti-Ly6G antibody (specific for mouse neutrophils) was administered intravenously (IV) to determine the location and concentration of neutrophils in mice. IV injection of LPS subjected mice to a model of mild ARDS. Accumulation of anti-Ly6G antibody in the lungs was dramatically affected by IV LPS, with 20.81% of injected antibody adhering in LPS-injured lungs, compared to 2.82% of injected antibody in naïve control lungs (Figure 1B). Agreeing with previous studies addressing the role of neutrophils in systemic inflammation, biodistributions of anti-Ly6G antibody indicated that systemic LPS injury profoundly increased the concentration of neutrophils in the lungs.^6,10,14,18^

Single cell suspensions prepared from mouse lungs were probed by flow cytometry to further characterize pulmonary neutrophils in naïve mice and in mice following LPS-induced inflammation. To identify intravascular populations of leukocytes, mice received IV fluorescent CD45 antibody five minutes prior to sacrifice. Single cell suspensions prepared from IV CD45-stained lungs were then stained with anti-Ly6G antibody to identify neutrophils. A second stain of single cell suspensions with CD45 antibody indicated the total population of leukocytes in the lungs, distinct from the intravascular population indicated by IV CD45.

Flow cytometry showed greater concentrations of neutrophils in LPS-injured lungs, compared to naïve lungs (Figure 1C, counts above horizontal threshold indicate positive staining for neutrophils, Figure 1D, rightmost peak indicates positive staining for neutrophils). Comparison of Ly6G stain to total CD45-positive cells indicated 53.5% of leukocytes in the lungs were Ly6G-positive after LPS injury, compared to 5.6% in the naïve control (Figure 1D, center panel). Comparison of Ly6G stain to IV CD45 stain indicated that the majority of neutrophils were intravascular, in both naïve and LPS-injured mice. In naïve mice, 85.3% of neutrophils were intravascular and in LPS-injured mice, 96.2% of neutrophils were intravascular (Figure 1D, right panel). The presence of large populations of intravascular neutrophils following inflammatory injury is consistent with previously published observations.^6,10,14,18,33^

Histological analysis confirmed results obtained with flow cytometry and radiolabeled anti-Ly6G biodistributions. Staining of lung sections indicated increased concentration of neutrophils in the lungs following IV LPS injury (Figure 1E, left panels). Co-registration of neutrophil staining with tissue autofluorescence (indicating tissue architecture) broadly supported the finding that pulmonary neutrophils reside in the vasculature (Figure 1E, right panels).

Previous work has traced the neutrophil response to bacteria in the lungs, determining that pulmonary neutrophils pursue and engulf active bacteria following either intravenous infection or infection of the airspace in the lungs.^18,34,35^ We injected heat-inactivated, oxidized, and fixed *E. coli* in naïve and IV-LPS-injured mice. With the bacteria stripped of their functional behavior, *E. coli* did not accumulate in the lungs of naïve control mice (1.47% of initial dose in the lungs, blue bars in Figure 1F). However, pre-treatment with LPS to recapitulate the inflammatory response to infection led to enhanced accumulation of the deactivated *E. coli* in the lungs (7.69% of initial dose in the lungs, red bars in Figure 1F). With *E. coli* structure maintained but *E. coli* function removed, the inactivated bacteria were taken up more avidly in lungs primed by an inflammatory injury.

## Injury-Specific Uptake of Nanoparticles in Pulmonary Marginated Neutrophils

In order to identify nanostructural parameters that correlate with nanoparticle uptake in inflamed lungs, we conducted an *in vivo* screen of a diverse array of nanoparticle drug carriers. The screen was based on the method used above for tracing inactivated bacteria: inject radiolabeled nanoparticles into mice and measure biodistributions, comparing naïve with IV-LPS mice. To validate that the radiotracing screen would measure uptake in pulmonary marginated neutrophils, we more fully characterized the *in vivo* behavior of two early hits in the screen.

Lysozyme-dextran nanogels (LDNGs, NGs) and poly(ethylene)glycol (PEG)- crosslinked albumin NPs have been characterized as targeted drug delivery agents in previous work.^36-38^ Here, LDNGs (136.4±3.6 nm diameter, 0.10±0.02 PDI, Supplementary Figure 1A) and PEG-crosslinked human albumin NPs (317.8±3.6 nm diameter, 0.14±0.05 PDI, Supplementary Figure 1B) were administered in naïve and IV-LPS-injured mice. Neither NP was functionalized with antibodies or other affinity tags. The protein component of each particle was labeled with ^125^I for tracing in biodistributions, and assessed 30 minutes after IV administration of NPs. Both absolute LDNG lung uptake and ratio of lung uptake to liver uptake registered a ∼25-fold increase between naïve control and LPS-injured animals (Figure 2A, Supplementary Table 1). Specificity for LPS-injured lungs was recapitulated with PEG-crosslinked human albumin NPs. Albumin NPs accumulated in naïve lungs at 6.34% injected dose per gram organ weight (%ID/g), and in LPS-injured lungs at 87.62 %ID/g, accounting for a 14-fold increase in lung uptake after intravenous LPS insult (Figure 2B, Supplementary Table 1).

**Figure 2.**
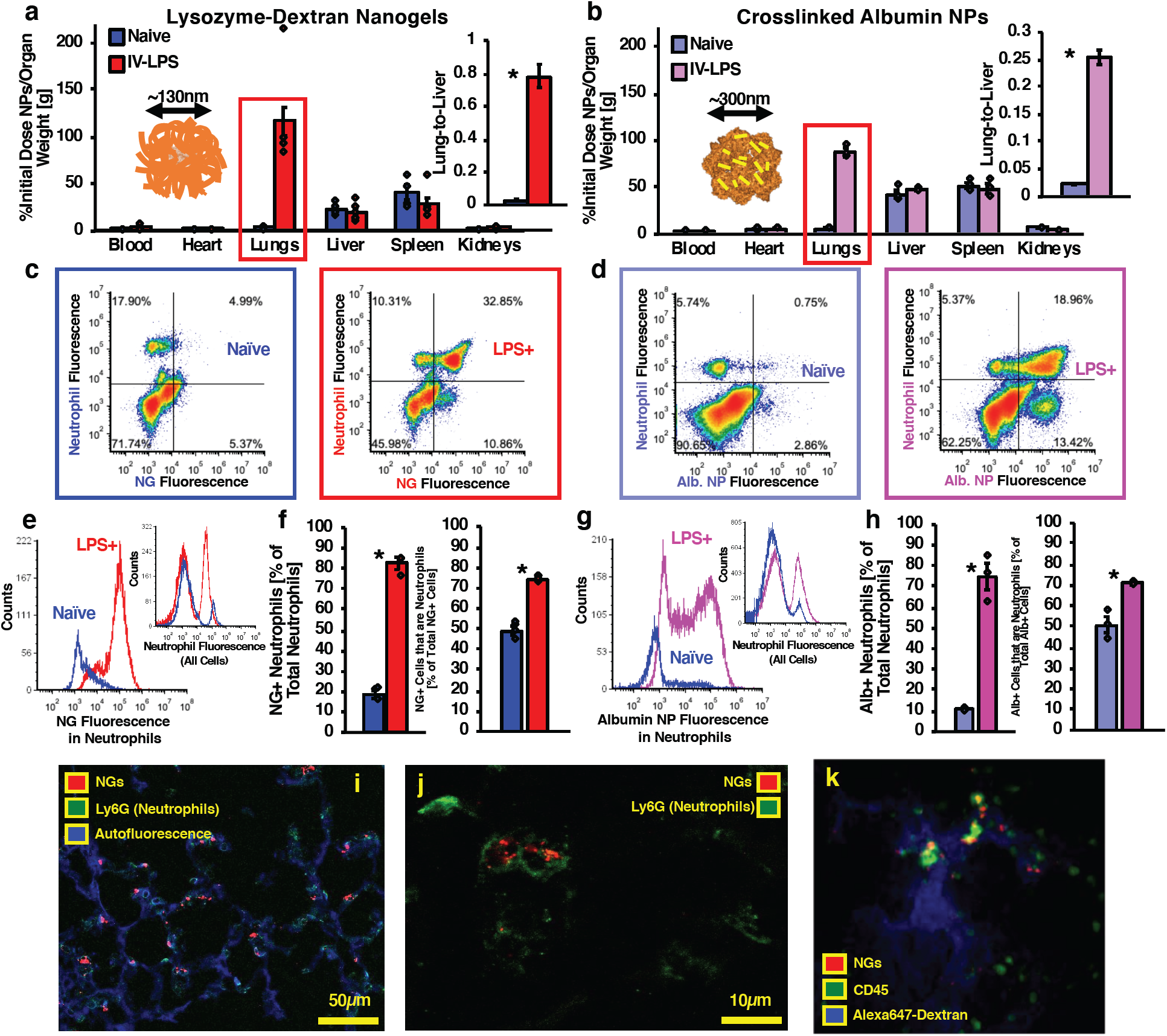
Lysozyme-Dextran Nanogels and Crosslinked Albumin Nanoparticles Accumulate in Marginated Neutrophils in Inflamed Lungs. (a) Biodistributions of lysozyme-dextran nanogels (LDNGs) in naïve (n=4) and IV-LPS-injured (n=8) mice (*red box = p*<.*001, * = p*<.*01*). (b) Biodistributions of PEG-NHS crosslinked human albumin nanoparticles (albumin NPs) in naïve (n=3) and IV-LPS-injured (n=3) mice (*red box = p*<.*001, * = p*<.*01*). (c-h) Flow cytometric characterization of single cell suspensions prepared from naïve and IV-LPS-injured injured mouse lungs. (c-d) Vertical axis indicates Ly6G staining (APC signal) and horizontal axis indicates signal from fluorescent LDNGs (c) or fluorescent albumin NPs (d). (e/g) LDNG/albumin NP fluorescent signal from neutrophils in IV-LPS-injured mouse lungs (red/pink), compared to naïve lungs (blue) (inset: Flow cytometry data verifying increased neutrophil concentration in IV-LPS-injured mouse lungs (red/pink). (f/h) Fraction of neutrophils positive for LDNGs (f) or albumin NPs (h) in naïve (blue, n=3) or IV-LPS-injured (red/pink, n=3) lungs and fraction of LDNG-positive (red, f) or albumin NP-positive (pink, h) cells that are neutrophils (*\* = p*<.*01*). (i-j) Fluorescence micrographs indicating association of LDNGs (red) with neutrophils (green, Ly6G stain) in IV-LPS-inflamed lungs (blue, tissue autofluorescence). (k) Single frame from real-time intravital imaging of LDNG (red) uptake in leukocytes (green) in IV-LPS-inflamed pulmonary vasculature (blue, Alexa Fluor 647-dextran).

Single cell suspensions were prepared from lungs after administration of fluorescent LDNGs or PEG-crosslinked albumin NPs. Flow cytometric analysis of cells prepared from lungs after NP administration enabled identification of cell types with which NPs associated. Firstly, the total number of cells containing LDNGs or albumin NPs increased between naïve and LPS-injured lungs. In naïve control lungs, 2.2% of cells were positive for LDNGs and 4.4% of cells were positive for albumin NPs. In LPS-injured lungs, 37.6% of cells were positive for LDNGs and 31.3% of cells were positive for albumin NPs (Supplementary Figure 2A, Supplementary Figure 2B, Supplementary Figure 3A, Supplementary Figure 3B).

Ly6G stain for neutrophils indicated that the bulk of LDNG and albumin NP accumulation in LPS-injured lungs could be accounted for by uptake in neutrophils. In Figure 2C and 2D, counts above the horizontal threshold indicate neutrophils and counts to the right of the vertical threshold indicate cells containing LDNGs (Figure 2C) or albumin NPs (Figure 2D). In IV-LPS-injured lungs, LDNG and albumin NP uptake was dominated by neutrophils (Figure 2C, Figure 2D, upper right quadrants indicate NP-positive neutrophils). In LPS-injured lungs, the majority of neutrophils, >70% of cells, contained significant quantities of nanoparticles, compared to <20% in naïve lungs. Likewise, the majority of nanoparticle uptake in the lungs (>70%) was accounted for by nanoparticle uptake in neutrophils (Figure 2E, 2F, 2G, 2H, Supplementary Table 2).

For NP uptake not accounted for by neutrophils, CD45 staining indicated that the remaining NP uptake was attributable to other leukocytes. Co-localization of albumin NP fluorescence with CD45 stain showed that 91.9% of albumin NP uptake was localized to leukocytes in naïve lungs and 97.8% of albumin NP uptake was localized to leukocytes in injured lungs (Supplementary Figure 3C, Supplementary Figure 3D).

For LDNGs, localization to neutrophils in injured lungs was confirmed via histology. Ly6G staining of LPS-injured lung sections confirmed colocalization of fluorescent nanogels with neutrophils in the lung vasculature (Figure 2I). Slices in confocal images of lung sections indicated that LDNGs were inside neutrophils (Figure 2J). Intravital imaging of injured lungs allowed real-time visualization of LDNG uptake in leukocytes in injured lungs. LDNG fluorescent signal accumulated over 30 minutes and reliably colocalized with CD45 staining for leukocytes (Figure 2K, Supplementary Movie 1).

LDNG pharmacokinetics were evaluated in naïve and IV-LPS-injured mice (Supplementary Figure 4). In both naïve and injured mice, bare LDNGs were rapidly cleared from the blood with a distribution half-life of ∼3 minutes. In naïve mice, transient retention of LDNGs in the lungs (25.91 %ID/g at five minutes after injection) leveled off over one hour. In IV-LPS-treated mice, LDNG concentration in the lungs reached a peak value at 30 minutes after injection, as measured either by absolute levels of lung uptake or by lungs:blood localization ratio.

LDNG biodistributions were also assessed in mice undergoing alternative forms of LPS-induced inflammation. Intratracheal (IT) instillation of LPS led to concentration of LDNGs in the lungs at 81.31 %ID/g. Liver and spleen LDNG uptake was also reduced following IT LPS injury, leading to a 45-fold increase in the lungs:liver LDNG localization ratio induced by IT LPS injury (Supplementary Figure 5). As with IV LPS injury, IT LPS administration leads to neutrophil-mediated vascular injury focused in the lungs.^14^

Mice were administered LPS via footpad injection to provide a model of systemic inflammation originating in lymphatic drainage.^39^ LDNG uptake in the lungs and in the legs was enhanced by footpad LPS administration. At 6 hours after footpad LPS administration, LDNGs concentrated in the lungs at 59.29 %ID/g, an 11-fold increase over naïve. At 24 hours, LDNGs concentrated in the lungs at 202.64 %ID/g (Supplementary Figure 6A). Total LDNG accumulation in the legs accounted for 0.85% of initial dose (%ID) in naïve mice, 2.65 %ID in mice 6 hours after footpad LPS injection, and 8.34 %ID at 24 hours after footpad injection (Supplementary Figure 6B), indicating LDNGs can concentrate in inflamed vasculature outside the lungs.

Previous work has indicated that NPs based on denatured albumin accumulate in neutrophils in inflamed lungs and at sites of acute vascular injury, whereas NPs coated with native albumin do not.^26,27^ We have characterized lysozyme-dextran nanogels and crosslinked human albumin NPs with circular dichroism (CD) spectroscopy to compare secondary structure of proteins in the NPs to secondary structure of the native component proteins (Supplementary Figure 7A-B). Identical CD spectra were recorded for LDNGs vs. lysozyme and for albumin NPs vs. human albumin. Deconvolution of the CD spectra via neural network algorithm trained against a library of CD spectra for known structures verified that secondary structure composition of lysozyme and albumin was unchanged by incorporation of the proteins in the NPs.^40^

Free protein and protein NPs were also probed with 8-anilino-1-naphthalenesulfonic acid (ANSA), previously established as a tool for determining the extent to which hydrophobic domains are exposed on proteins.^41^ Consistent with known structures of the two proteins, ANSA staining indicated few available hydrophobic domains on lysozyme and substantial hydrophobic exposure on albumin (Supplementary Figure 7C-D, blue curves). LDNGs had increased hydrophobic accessibility vs. native lysozyme whereas albumin NPs had reduced hydrophobic accessibility compared with native albumin. Therefore, our data indicate that lysozyme and albumin are not denatured in LDNGs and albumin NPs, but the NPs composed of the two proteins present a balance of hydrophobic and hydrophilic surfaces differing from the native proteins.

## In Vivo Screen of Diverse Protein Nanoparticle Structures in Acutely Inflamed Lungs

The previous section demonstrates that two different nanoparticles based on protein, shown not to be denatured in CD spectroscopy studies, have uptake in LPS-inflamed lungs driven by uptake in marginated neutrophils. We next undertook a broader study considering how aspects of NP structure including size, composition, surface chemistry, and structural organization impact NP uptake in LPS-injured lungs. As examples of different types of protein NPs, variants of LDNGs (representing NPs based on hydrophobic interactions between proteins), crosslinked protein NPs, and NPs based on electrostatic interactions between proteins were traced in naïve control and IV-LPS-injured mice. As examples of NPs based on site-specific protein interactions (rather than site-indiscriminate interactions leading to crosslinking, gelation, or charge-based protein NPs), we also traced viruses and ferritin nanocages in naïve and LPS-treated mice. Liposomes and polystyrene NPs were studied as examples of lipid and polymeric nanostructures.

### Nanoparticles Based on Hydrophobic Protein Interactions

LDNG size was varied by modifying lysozyme-dextran composition of the NPs and pH at which particles were formed.^42^ LDNGs of ∼75nm (73.2±1.3 nm, PDI 0.18±0.05), ∼200nm (199.4±1.8 nm, PDI 0.11±0.01), and ∼275nm (274.5±6.4 nm, PDI 0.16±0.06) diameter were traced in naïve control and IV-LPS-injured mice, adding to data obtained for 130nm LDNGs above (Figure 3A, Supplementary Figure 1A, Supplementary Figure 8). As with data for 130nm LDNGs reported in Figure 2, all sizes of LDNGs accumulated in LPS-injured lungs at higher concentrations than in naïve lungs, with accumulation in injured lungs reaching ∼20% of initial dose for all types of LDNGs (Supplementary Table 3). *Variations in size and composition of LDNGs therefore did not affect LDNG specificity for LPS-injured lungs*.

**Figure 3.**
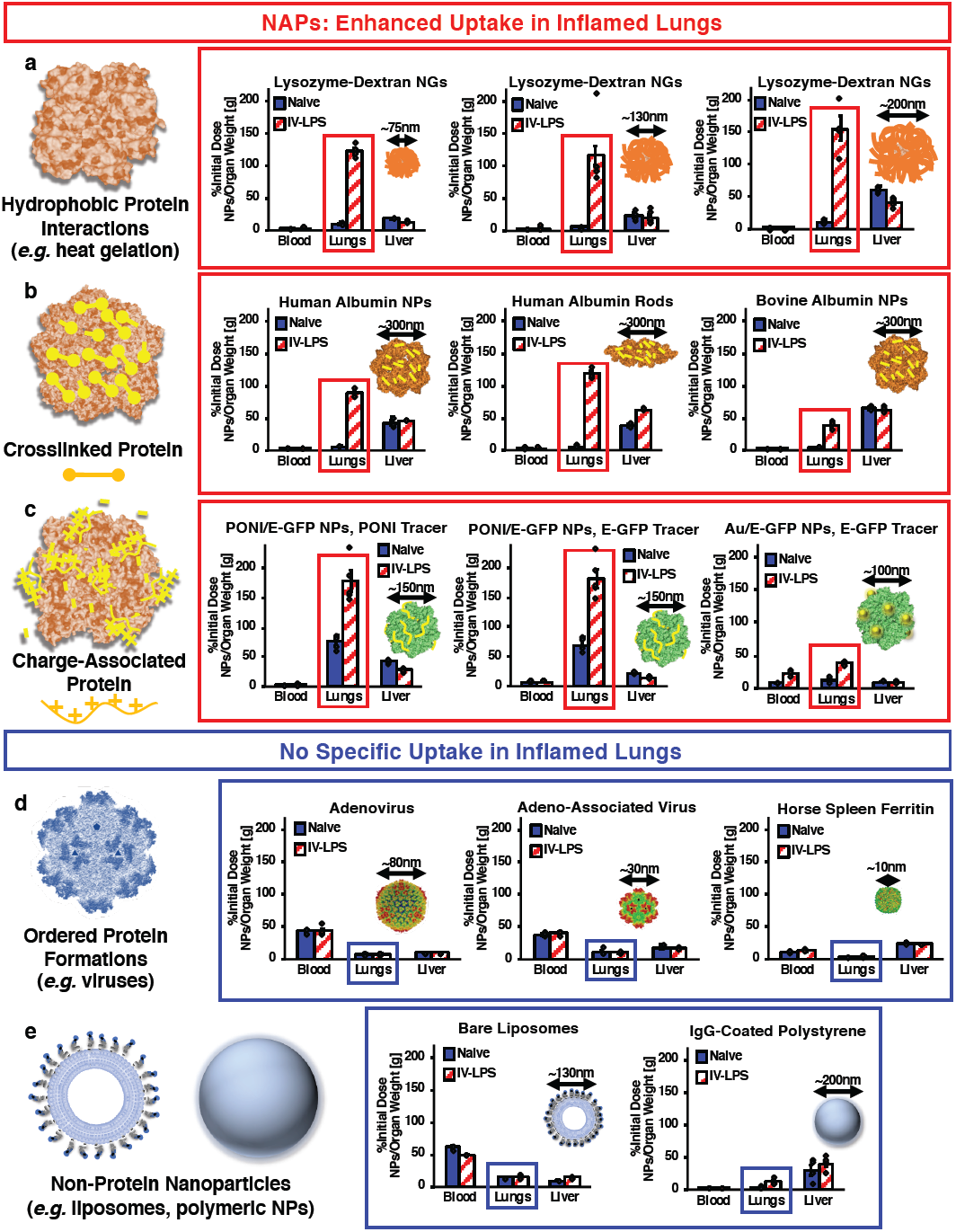
Screen of Diverse Nanoparticle Biodistributions in Naïve and IV-LPS-Inflamed Lungs. (a-c) Nanoparticles with agglutinated protein (NAPs) accumulate in acutely inflamed lungs. (a) Biodistributions of variant lysozyme-dextran nanogels indicating uptake of 75nm lysozyme-dextran nanogels (n=4 IV-LPS, n=4 naïve) and 200nm lysozyme-dextran nanogels (n=5 IV-LPS, n=5 naïve) in LPS-injured lungs, but not naïve lungs. Data for 130nm lysozyme-dextran nanogels is identical to that presented in Figure 2a. (b) Biodistributions of variant crosslinked albumin nanoparticles indicating uptake of albumin nanorods (n=3 IV-LPS, n=3 naïve) and bovine albumin nanoparticles (n=3 IV-LPS, n=3 naïve) in LPS-injured, but not naïve lungs. Data for human albumin nanoparticles is identical to that presented in Figure 2b. (c) Biodistributions of charge-agglutinated protein nanoparticles, indicating uptake of particles comprised of glutamate-tagged green fluorescent protein (E-GFP) and guanidine-tagged poly(oxanorborneneimide) (PONI) or particles comprised of E-GFP and guanidine-tagged gold nanoparticles (Au) in LPS-injured (PONI: n=5, Au: n=3), but not naïve (PONI: n=4, Au: n=3) lungs. PONI/E-GFP data reflects tracing of both ^131^I-labeled PONI and ^125^I-labeled E-GFP. (d-e) Nanoparticles based on highly symmetric supramolecular organization of protein or nanoparticles not based on protein do not have tropism for inflamed lungs. (d) Biodistributions of adenovirus (n=5 IV-LPS, n=5 naive), adeno-associated virus (n=3 IV-LPS, n=3 naïve), and ferritin nanocages (n=5 IV-LPS, n=5 naïve) indicating no specificity for LPS-injured vs. naïve lungs. (e) Biodistributions of bare liposomes (n=4 IV-LPS, n=4 naïve) indicating no specificity for LPS-injured vs. naïve lungs. Biodistributions of IgG-coated polystyrene nanoparticles indicating low levels of uptake in both naïve (n=4) and LPS-injured (n=4) lungs. (For statistical comparison of NP uptake in inflamed vs. naïve lungs: *red box = p*<.*001, blue box = no significance at p*<.*001 level*).

### Nanoparticles Based on Protein Crosslinking

Expanding on data with PEG-NHS ester-crosslinked human serum albumin particles, we varied the geometry and protein composition of NPs based on PEG-NHS protein crosslinking. Human serum albumin nanorods (aspect ratio 3:1), bovine serum albumin NPs (317.3±38.5 nm, PDI 0.17±0.04), human hemoglobin NPs (328.1±16.1 nm, PDI 0.08±0.01), human transferrin NPs (345.2±10.2 nm, PDI 0.12±0.004), and chicken lysozyme NPs (298.6±12.4 nm, PDI 0.06±0.01) were traced in naïve and IV LPS-injured mice (Figure 3B, Supplementary Figure 1B, Supplementary Figure 9). With the exception of crosslinked lysozyme NPs, all of the tested formulations had clear specificity for acutely inflamed lungs over naïve lungs (Supplementary Table 4). Lysozyme NPs accumulated in naïve lungs at a uniquely high concentration of 137.47 %ID/g, compared to 170.92 %ID/g in inflamed lungs. Degree of uptake in injured lungs, along with injured vs. naïve contrast, did vary with protein NP composition. However, *acute inflammatory injury resulted in a minimum three-fold increase in lung uptake for all examined crosslinked protein NPs, excluding crosslinked lysozyme, which still accumulated in injured lungs at a high concentration (25.64% of initial dose).*

### Nanoparticles Based on Electrostatic Protein Interactions

We traced recently-developed poly(glutamate) tagged green fluorescent protein (E-GFP) NPs, representing a third class of protein NP based on electrostatic interactions between proteins and carrier polymer or metallic particles.^43^ Negatively-charged E-GFP was paired to arginine-presenting gold nanoparticles (89.0±1.6 nm diameter, PDI 0.14±0.04) or to poly(oxanorborneneimide) (PONI) functionalized with guanidino and tyrosyl side chains (158.9±6.2 nm diameter, PDI 0.17±0.03) (Supplementary Figure 1D). For biodistribution experiments with PONI/E-GFP hybrid NPs, tyrosine-bearing PONI was labeled with ^131^I and E-GFP was labeled with ^125^I, allowing simultaneous tracing of each component of the hybrid NPs. The two E-GFP NPs, with structure based on charge interactions, had specificity for IV LPS-injured lungs. Comparing uptake in LPS-injured lungs to naïve lungs, we observe an LPS:naïve ratio of 2.37 for PONI/E-GFP NPs as traced by the PONI component, 2.57 for PONI/E-GFP NPs as traced by the E-GFP component, and 2.79 for Au/E-GFP NPs (Figure 3C, Supplementary Figure 10). PONI/E-GFP particles, specifically, accumulated in LPS-injured lungs at 26.77% initial dose as measured by PONI tracing and 27.24% initial dose as measured by GFP tracing, indicating effective co-delivery in the inflamed organ. *Acute inflammatory injury therefore resulted in a two-to three-fold increase in pulmonary uptake of NPs constructed via electrostatic protein interactions.*

### Nanoparticles Based on Symmetric Protein Organization

Adeno-associated virus (AAV), adenovirus, and horse spleen ferritin nanocages were employed as examples of protein-based NPs with highly symmetrical structure (See Supplementary Figure 1D for DLS confirmation of structure).^44-46^ For each of these highly ordered protein NPs, IV LPS injury had no significant effect on biodistribution and levels of uptake in the injured lungs were minimal (Figure 3D, Supplementary Figure 11, Supplementary Table 5). Therefore, *highly ordered protein NPs traced in our studies did not have tropism for the lungs after acute inflammatory injury*.

### Non-Protein Nanoparticles

Liposomes and polystyrene NPs were studied as example NPs that are not structurally based on proteins. DOTA chelate-containing lipids were incorporated into bare liposomes, allowing labeling with ^111^In tracer for biodistribution studies.

Carboxylate polystyrene NPs were coupled to trace amounts of ^125^I-labeled IgG via EDCI-mediated carboxy-amine coupling. Liposomes had a diameter of 103.6±8.7 nm (PDI 0.09±0.01) and IgG-polystyrene NPs had a diameter of 230.5±2.8 nm (PDI 0.14±0.01) (Supplementary Figure 1C-D). Liposomes accumulated in inflamed lungs at a concentration of 16.89 %ID/g, accounting for no significant change against naïve lungs. LPS injury actually induced a reduction in the lungs:liver metric, from 0.20 for naïve mice to 0.15 for LPS-injured mice. Polystyrene NPs accumulated in inflamed lungs at 11.67 %ID/g (1.75% initial dose), so IV LPS injury did in fact induce increased levels of NP uptake in the lungs, from a concentration of 2.40 %ID/g in the naïve lungs (Figure 3E, Supplementary Figure 12). However, *neither bare liposomes nor polystyrene NPs were drawn to LPS-injured lungs in significant concentrations*.

### Isolated Proteins

Significantly, isolated proteins did not home to LPS-inflamed lungs themselves. We traced radiolabeled albumin, lysozyme, and transferrin in naïve control and IV LPS-injured mice (Supplementary Figure 13, Supplementary Table 6). *In injured mice, albumin, lysozyme, and transferrin localized to the lungs at low concentrations and no significant differences were recorded when comparing naïve to LPS-injured lung uptake*.

The data presented in figure 3 and supplementary figures 8-13 indicate that a variety of protein-based nanostructures have tropism for acute inflammatory injury in the lungs. NPs based on agglutination of proteins in non-site-specific interactions (NAPs, Figure 3A-C, Supplementary Figures 8-10) all exhibited either significant increases in lung uptake after LPS injury or high levels of lung uptake in both naïve control and LPS-injured animals. Nanostructures based on highly symmetrical protein organization had no specific tropism for inflamed lungs (Figure 3D). Representative nanostructures not based on proteins, bare liposomes and polystyrene beads, did not home to inflamed lungs (Figure 3E).

## Engineering of Liposomes for Structure-Based Neutrophil Tropism in Acutely Injured Lungs

We next engineered NAPs from liposomes, a nanoparticle shown above to have no intrinsic neutrophil tropism. Our methods for engineering NAP-like liposomes serve to validate the finding that supramolecular organization of protein in nanoparticles predicts neutrophil tropism.

Liposomes were functionalized with rat IgG conjugated via SATA-maleimide chemistry (SATA-IgG liposomes) or via recently demonstrated copper-free click chemistry methods.^47^ Briefly, click chemistry methods entailed NHS-ester conjugation of an excess of strained alkyne (dibenzocyclooctyne, DBCO) to IgG, followed by reaction of the DBCO-functionalized IgG with liposomes containing PEG-azide-terminated lipids (DBCO-IgG liposomes, Figure 4A). DBCO-IgG liposomes had a diameter of 128.3±4.3 nm and a PDI of 0.17±0.03 and SATA-IgG liposomes had a diameter of 178.8±6.9 nm and a PDI of 0.23±0.03 (Supplementary Figure 1C).

**Figure 4.**
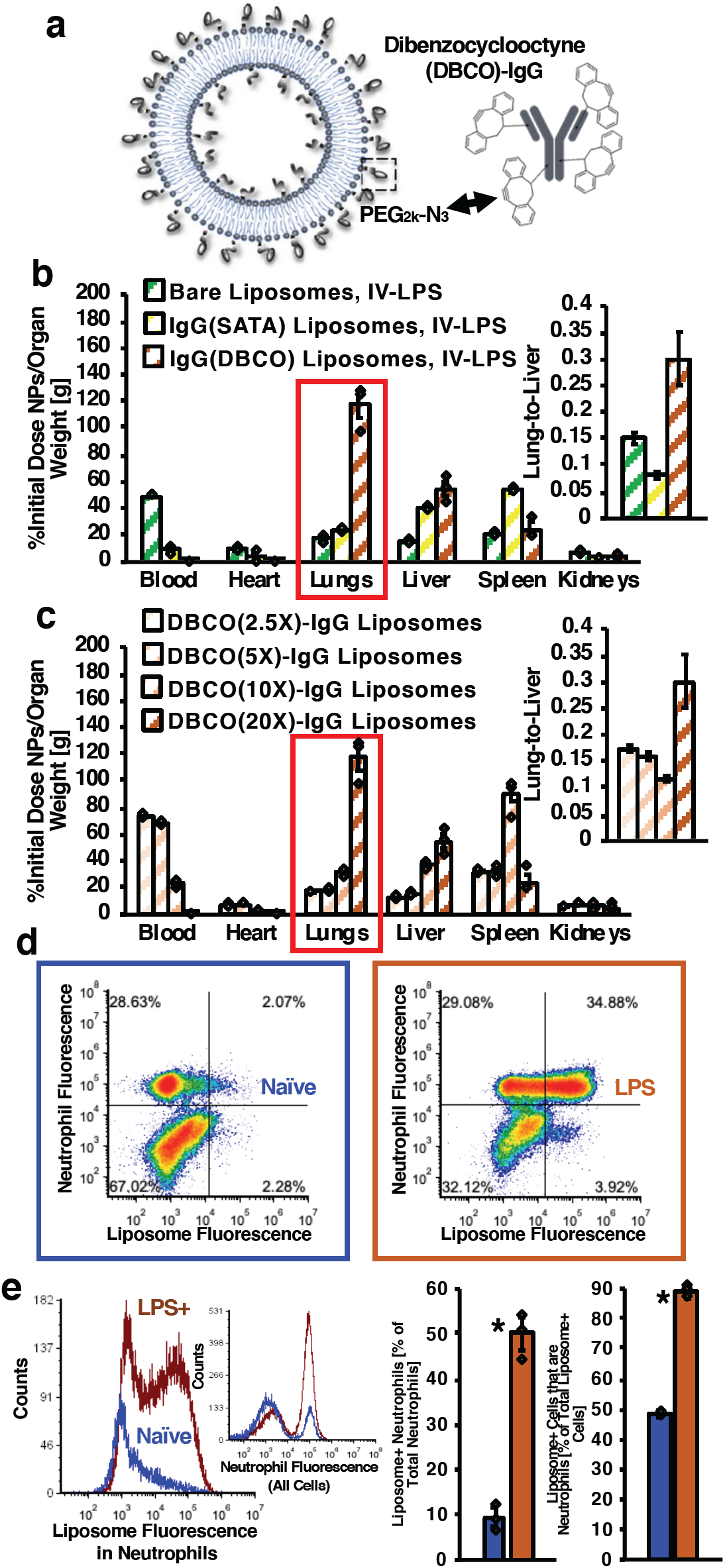
Engineering of Liposome Surface Chemistry to Confer NAP-like Behavior in LPS-Inflamed Lungs. (a) Schematic of antibody-coated liposomes prepared via copper-free click reaction of azide-functionalized liposomes with dibenzocyclooctyne (DBCO)- functionalized IgG. (b) Biodistributions in IV-LPS-injured mice for bare liposomes (green stripes), liposomes conjugated to IgG via SATA-maleimide chemistry (yellow stripes), and liposomes conjugated to IgG via DBCO-azide chemistry (brown stripes). (c) Biodistributions in IV-LPS-injured mice for azide-functionalized liposomes conjugated to IgG loaded with 2.5, 5, 10, and 20 DBCO molecules per IgG (bars further to the right correspond to more DBCO per IgG). (d) Mouse lungs flow cytometry data indicating Ly6G anti-neutrophil staining density vs. levels of DBCO(20X)-IgG liposome uptake. (e) Flow cytometry data verifying increased DBCO(20X)-IgG liposome uptake in and specificity for neutrophils following LPS insult (Inset: Verification of increased concentration of neutrophils in the lungs following LPS) (*red box = p*<.*001, * = p*<.*01*).

In mice subjected to IV-LPS, SATA-IgG liposomes accumulated in the lungs at a concentration of 22.26 %ID/g (Figure 4B, yellow bars). DBCO-IgG liposomes, by contrast, concentrated in the lungs at 117.16 %ID/g, corresponding to 17.57% of initial dose and roughly matching the accumulation of 130nm LDNGs in the inflamed lungs (Figure 4B, brown bars). For comparison, bare liposomes, as in Figure 3E, concentrated in the inflamed lungs at 16.89 %ID/g (Figure 4B, green bars). For DBCO-IgG liposomes, the inflamed vs. naïve lung uptake accounted for a twelve-fold change. DBCO-IgG liposomes specifically accumulated in injured lungs, whereas SATA-IgG liposomes and bare liposomes did not (Supplementary Figure 14, Supplementary Table 7).

IT LPS instillation also led to elevated concentrations of DBCO-IgG liposomes in the lungs. Biodistributions of the DBCO-IgG liposomes indicated a pulmonary concentration of 145.89 %ID/g at 1 hour after IT LPS, 160.13 %ID/g at 2 hours after IT LPS, and 127.78 %ID/g at 6 hours after IT LPS (Supplementary Figure 15). Even at early time points after direct pulmonary LPS insult, DBCO-IgG liposomes accumulated in the inflamed lungs.

Results in Figure 4B were obtained by introducing a 20-fold molar excess of NHS-ester-DBCO to rat IgG before DBCO-IgG conjugation to liposomes (DBCO(20X)- IgG liposomes). Optical density quantification of DBCO indicated ∼14 DBCO per IgG following reaction of DBCO and IgG at 20:1 molar ratio (Supplementary Figure 16). To test the hypothesis that DBCO functions as a tag that modifies DBCO-IgG liposomes for neutrophil affinity in settings of inflammation, we varied the concentration of DBCO on IgG prepared for conjugation to azide liposomes. DBCO was added to IgG at 10-fold, five-fold, and 2.5-fold molar excesses. A 10-fold molar excess resulted in ∼6 DBCO per IgG, a 5-fold molar excess resulted in ∼3 DBCO per IgG, and a 2.5-fold molar excess resulted in ∼2 DBCO per IgG (Supplementary Figure 16). IgG with different DBCO loading concentrations was conjugated to azide liposomes. DBCO-IgG liposomes had similar sizes across all DBCO concentrations (Supplementary Figure 1C), with diameters of ∼130 nm and PDIs < 0.20.

The different types of DBCO-IgG liposomes were each traced in IV-LPS injured mice. Titrating the quantity of DBCO on DBCO-IgG liposomes indicated that liposome accumulation in the lungs of injured mice was dependent on DBCO concentration on the liposome surface. Concentration of DBCO-IgG liposomes in inflamed lungs attenuated with decreasing DBCO concentration on IgG (Supplementary Table 8, Figure 4C). Therefore, only IgG with high concentrations of DBCO served as a tag for modifying the surface of liposomes for specificity to pulmonary injury.

Flow cytometry verified the specificity of DBCO-IgG liposomes for neutrophils in injured lungs (Figure 4D-E). As with LDNGs and albumin NPs in Figure 2C-H, single cell suspensions were prepared from LPS-inflamed and naïve control lungs after circulation of fluorescent DBCO-IgG liposomes. Confirming the results of biodistribution studies, 4.90% of cells were liposome-positive in naïve lungs, compared to 33.92% of all cells in LPS-inflamed lungs (Supplementary Figure 17A-B).

DBCO-IgG liposomes predominantly accumulated in pulmonary neutrophils after IV LPS. There were more neutrophils in the injured lungs and a greater fraction of neutrophils took up DBCO-IgG liposomes in the injured lungs, as compared to the naïve control (Figure 4D-E). Approximately one half of neutrophils in IV LPS-injured lungs contained liposomes. DBCO-IgG liposomes were also highly specific for neutrophils in inflamed lungs, with ∼90% of liposome-positive cells in the injured lungs being neutrophils (Supplementary Table 9). The remaining DBCO-IgG liposome uptake in the lungs was accounted for by other CD45-positive cells (Supplementary Figure 17C-E). 99.0% of liposome uptake colocalized with CD45-positive cells in LPS-injured lungs and 98.7% of liposome uptake in the naïve lungs was associated with CD45-positive cells. Accordingly, less than 1% of liposome uptake was associated with endothelial cells (Supplementary Figure 17F-G).

DBCO(20X)-IgG itself did not have specificity for inflamed lungs (Supplementary Figure 18). Uptake of DBCO(20X)-IgG in naïve and injured lungs was statistically identical and the biodistribution of the modified IgG resembled published results with unmodified IgG.^48^ These results verify that DBCO-IgG modifies the structure of immunoliposomes, but does not function as a standard affinity tag by acting as a surface motif with intrinsic affinity for neutrophils.

Indeed, CD spectroscopic and ANSA structural characterization of DBCO-modified IgG and DBCO-IgG liposomes resembled results obtained for LDNGs and crosslinked albumin NPs. IgG secondary structure, as assessed by CD spectroscopy, was unchanged by DBCO modification (Supplementary Figure 19A). Deconvolution of CD spectra via neural network algorithm indicated identical structural compositions for DBCO(20X)-IgG, DBCO(10X)-IgG, DBCO(5X)-IgG, DBCO(2.5X)-IgG, and unmodified IgG, showing that IgG was not denatured by conjugation to DBCO. ANSA was used to probe accessible hydrophobic domains on DBCO(20X)-IgG and DBCO(20X)-IgG liposomes (Supplementary Figure 19B). ANSA fluorescence indicated more hydrophobic domains available on DBCO(20X)-IgG liposomes than on DBCO(20X)-IgG itself, resembling results for lysozyme and LDNGs.

Therefore, addition of a hydrophobic moiety to protein on the surface of liposomes led to uptake of the liposomes in pulmonary marginated neutrophils after inflammatory insult. This result indicates that hydrophobic interactions between proteins on the surface of functionalized liposomes, like the protein interactions in NAPs, predict liposome tropism for marginated neutrophils in inflamed lungs.

## Principal Component and Linear Discriminant Analysis of Nanoparticle Biodistributions

Including NPs from our four classes of protein-based NPs, two non-protein NPs (bare liposomes and polystyrene NPs), and five types of IgG-coated liposomes, we traced 23 nanoparticles in naïve and inflamed mice. Direct assessment of naïve-to-inflamed shifts in lung uptake led us to identify 13 NAPs with specificity for inflamed lungs. To verify this assessment and derive additional patterns in the broader data set, we undertook linear discriminant and principal components analyses of the biodistribution data for our 23 nanoparticles, along with three isolated proteins.

Grouping the 23 nanoparticles and three proteins according to the classes defined in Figure 3 and Supplementary Figures 8-13, we completed a linear discriminant analysis of the naïve-to-inflamed shift for particle retention in the lungs, blood, liver, and spleen (Supplementary Figure 20a). Data for particle uptake in each organ was normalized by subtracting and then dividing by the mean uptake over all particles. The first two eigenvectors, dominated by splenic uptake and a combination of liver and lung uptake, respectively, accounted for 96% of variation in the data. The resulting projection of the data along the first two linear discriminant analysis eigenvectors was analyzed by k-means clustering to confirm the classes of nanoparticle with specificity for the inflamed lungs (Supplementary Figure 20b). Indeed, division of the data into two clusters supported the delineation of the 13 nanoparticles with specificity for inflamed lungs. NAPs, nanoparticles based on protein gelation, crosslinking, and charge association, all aligned in one cluster.

As an exception, DBCO(20X)-IgG liposomes were considered as a unique class of particle and the linear discriminant analysis indicated that the inflammation-specific liposomes had *in vivo* behavior resembling that of LDNGs or PONI-GFP nanoparticles. This analysis of the liposome biodistributions supports the classification of DBCO(20X)- IgG liposomes as NAPs. IgG-coated polystyrene nanoparticles and DBCO(10X)-IgG liposomes were part of the k-means cluster without inflammation specificity, but data for these two particles resided close to the Voronoi boundary distinguishing the two clusters.

Principal component analysis comparing normalized nanoparticle uptake in inflamed lungs to normalized retention in liver, spleen, and blood provided a reductive metric to compare the distinct *in vivo* behavior of nanoparticles in the classes identified by linear discriminant analysis. Most variation in the biodistribution data was accounted for by an eigenvector closely aligned to variation in pulmonary uptake (Supplementary Figure 21a). Data was projected along that first eigenvector and magnitude of the projection was determined for each nanoparticle (Supplementary Figure 21b). First eigenvector projection values were then grouped according to the classes examined above via linear discriminant analysis. Only the classes in the inflammation-specific k-means cluster had positive average first eigenvector projections. All other particle classes had average first eigenvector projections indistinguishable from isolated protein (Supplementary Figure 21b).

Principal component and linear discriminant analyses of our compiled biodistributions confirmed; a) identification of NAPs as nanoparticles with distinct tropism for inflamed lungs and; b) alignment of DBCO(20X)-IgG liposome *in vivo* behavior with that of other NAPs.

## Imaging Lung Inflammation with NAPs

Computerized tomography (CT) imaging is a standard diagnostic tool for ARDS. CT images can identify the presence of edematous fluid in the lungs, but CT cannot distinguish between the two major types of pulmonary edema: non-inflammatory cardiogenic pulmonary edema (CPE) and ARDS-associated edema.^49^ We sought to use NAPs to distinguish inflammatory lung injury from CPE in diagnostic imaging experiments.

We induced CPE in mice via prolonged IV propranolol infusion.^50^ Edema was confirmed via CT imaging of inflated lungs *ex vivo* and *in situ*. Three-dimensional reconstructions of chest CT images were partitioned to distinguish airspace and low-density tissue, as in normal lungs (white, yellow, and light orange signal in Figure 5a), from high-density tissue and edema (red and black/transparent signal in Figure 5a).

**Figure 5.**
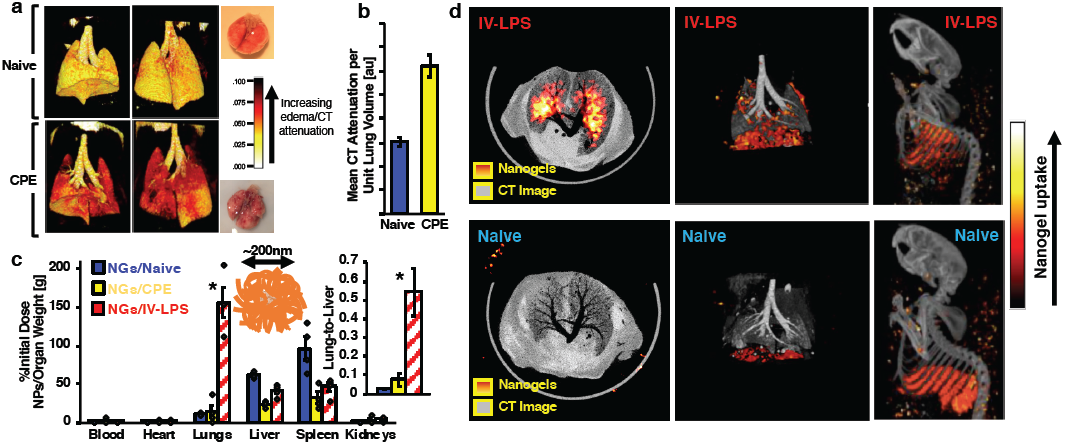
Specificity of NAPs for LPS-Inflamed Lungs vs. Edematous Lungs and SPECT Imaging of NAPs in LPS-Inflamed Lungs. (a) Three-dimensional reconstructions of chest CT data for naïve mouse lungs and lungs with cardiogenic pulmonary edema (CPE). White/yellow indicates lower attenuation, corresponding to airspace in healthy lungs. Red/dark background indicates higher attenuation, corresponding to fluid in the lungs. (b) Quantification of CT attenuation in naive (blue) and edematous (yellow) lungs. (c) Biodistributions of 200nm lysozyme-dextran nanogels in naïve mice (blue), mice treated with IV-LPS (red stripes), and mice subject to cardiogenic pulmonary edema (yellow). Naïve and IV-LPS data is identical to that presented in Supplementary Figure 8. (d) Co-registered CT (greyscale) and SPECT (red-yellow) images indicating ^111^In-labeled lysozyme-dextran nanogel uptake in naïve and IV-LPS injured mice. White/yellow indicates more nanogel uptake. Red/dark background indicates less nanogel uptake (*\* = p*<.*001*).

Quantification of CT attenuation and gaps in the reconstructed three-dimensional lung images indicated profuse edema in lungs afflicted with model cardiogenic pulmonary edema (Figure 5A-B, Supplementary Figure 20, Supplementary Movies 2 and 3). 200 nm LDNGs were traced in mice with induced cardiogenic pulmonary edema. LDNGs accumulated in the edematous lungs at 14.52 %ID/g concentration, statistically indistinguishable from lung uptake in naïve mice and an order of magnitude lower than the level of lung uptake in mice treated with IV LPS (Figure 5C).

Naïve and IV LPS-injured mice were dosed with LDNGs labeled with ^111^In via chelate conjugation to lysozyme. ^111^In uptake in naïve and LPS-injured lungs was visualized with *ex vivo* SPECT-CT imaging to indicate capacity of LDNGs for imaging-based diagnosis of inflammatory lung injury (Figure 5D). ^111^In signa was colocalized with anatomical CT images for reconstructions in Figure 5D. ^111^In SPECT signal was detectable in LPS-injured lungs, but ^111^In SPECT signal was at background level in naïve lungs (Supplementary Movies 4 and 5). Reduced SPECT signal in the liver of LPS-injured mice, in agreement with biodistribution data, was also evident in co-registration of SPECT imaging with full body skeletal CT imaging (Supplementary Movies 6 and 7).

Therefore, NAPs with tropism for marginated neutrophils have the ability to detect and assess ARDS-like inflammation via SPECT-CT imaging. Since those same NAPs do not accumulate in lungs afflicted with CPE, NAPs have potential for differential diagnosis of acute lung inflammation against CPE.

## NAP Accumulation in Inflamed Human Lungs Ex Vivo

In recent work, we demonstrated that human donor lungs rejected for transplant due to ARDS-like phenotypes can be perfused with nanoparticle solutions.^51^ These perfusion experiments evaluate the tendency of nanoparticles to distribute to human lungs *ex vivo*. We used this perfusion method to evaluate NAP retention in inflamed human lungs.

First, fluorescent LDNGs were added to single cell suspensions prepared from human lungs. 5 μg, 10 μg, or 50 μg of LDNGs were incubated with 6×10^5^ cells in suspension for 1 hour at room temperature. After three washes to remove unbound LDNGs, cells were stained for CD45 and analyzed with flow cytometry (Figure 7A-B). The majority of LDNG uptake in the single cell suspensions was attributable to CD45-positive cells. LDNGs accumulated in the human leukocytes, extracted from inflamed lungs, in a dose-dependent manner, with 35.1% of leukocytes containing LDNGs at a loading dose of 50 μg. Therefore, our prototype NAP was retained in leukocytes from human lungs.

To test LDNG tropism for inflamed intact human lungs, fluorescent or ^125^I-labeled LDNGs were infused via arterial catheter into *ex vivo* human lungs excluded from transplant. Immediately prior to LDNG administration, tissue dye was infused via the same arterial catheter to stain regions of the lungs directly perfused by the catheterized branch of the pulmonary artery (Figure 7C). After infusion of LDNGs, phosphate buffered saline infusion was used to rinse away unbound particles. Perfused regions of the lungs were dissected and divided into ∼1g segments, then sorted into regions deemed to have high, medium, or low levels of tissue dye staining. For lungs receiving fluorescent LDNGs, well-perfused and poorly-perfused regions were selected for sectioning and fluorescent imaging. Fluorescent signal from LDNGs was clearly detectable in sections of well-perfused tissue, but not poorly-perfused tissue (Figure 7D). In experiments with ^125^I-labeled LDNGs, ^131^I-labeled ferritin was concurrently infused (*i.e.* a mix of ferritin and LDNGs was infused) as an internal control particle shown to have no tropism for injured mouse lungs. With LDNGs and ferritin infused into the same lungs via the same branch of the pulmonary artery, LDNGs retained in the lungs at 52.15% initial dose and ferritin retained at 9.27% initial dose (Figure 7E). LDNG accumulation in human lungs was focused in regions of the lungs with high levels of perfusion stain, with concentrations of 4.66 %ID/g in the “high” perfusion regions, compared to 0.44 %ID/g in the “medium” perfusion regions. Ferritin accumulation was more diffuse, with 0.47 %ID/g in the “high” perfusion regions, compared to 0.35 %ID/g in the “medium” perfusion regions (Supplementary Figure 21).

LDNGs, a prototype NAP shown to home to neutrophils in acutely inflamed mouse lungs, specifically accumulated in perfused regions of inflamed human lungs, but ferritin nanocages, a particle with no tropism for neutrophils, concentrated at much lower levels in injured human lungs. Our data thus indicate that NAP tropism for neutrophils in inflamed mouse lungs may be recapitulated in human lungs.

## Therapeutic Effects of NAPs in Model ARDS

Previous studies indicate that nanoparticles can interfere with neutrophil adhesion in inflamed vasculature.^52^ We designed studies to evaluate whether or not NAPs mitigate the neutrophil-mediated effects of lung inflammation. Namely, we administered LDNGs, DBCO(20X)-IgG liposomes, or bare liposomes in mice subjected to model ARDS and determined whether or not the nanoparticles prevented lung edema induced by inflammation.

Mice were treated with nebulized LPS as a high-throughput model for severe ARDS. To evaluate physiological effects of the model injury, bronchoalveolar lavage (BAL) fluid was harvested from mice at 24 hours after exposure to LPS. In three separate experiments, nebulized LPS induced elevated concentrations of neutrophils, CD45-positive cells, and protein in the BAL fluid. In naïve mice, CD45-positive cells concentrated at 1.42×10^4^ cells per mL BAL and neutrophils concentrated at 1.11×10^4^ cells per mL BAL. After LPS injury, CD45-positive cells and neutrophils concentrated at 6.97×10^5^ and 6.96×10^5^ cells per mL BAL, respectively. In naïve mice, protein concentrated in the BAL fluid at 0.12 mg/mL and in LPS-injured mice, protein concentrated in the BAL at 0.36 mg/mL (Figure 6, white and grey bars). Vascular disruption after nebulized LPS treatment thus led to accumulation of protein-rich edema in the alveolar space.

**Figure 6.**
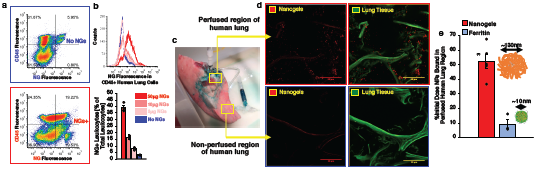
Uptake of NAPs in Ex Vivo Human Lungs. (a-b) Flow cytometry characterization of single cell suspensions prepared from donor human lungs rejected for transplant. (a) CD45 staining vs. lysozyme-dextran nanogel uptake in cells for cells without nanogels (blue) and cells incubated with nanogels (red, 50µg nanogels per 2×10^6^ cells). (b) Upper panel: Nanogel fluorescence in CD45-positive cells, after cell incubation with different quantities of nanogels. Lower panel: Percentage of leukocytes with nanogels after incubation of cells with different quantities of nanogels. (c) Photograph depicting cannulation of human lung for nanoparticle and tissue dye infusion, with green tissue dye indicating perfused regions of the lung. (d) Micrographs indicating fluorescent nanogel uptake in perfused and non-perfused regions of human lung. (e) Radiotracer-determined quantity of nanoparticle uptake in human lungs for lysozyme-dextran nanogels and ferritin simultaneously infused in the same lungs (*\* = p*<.*001*).

**Figure 7.**
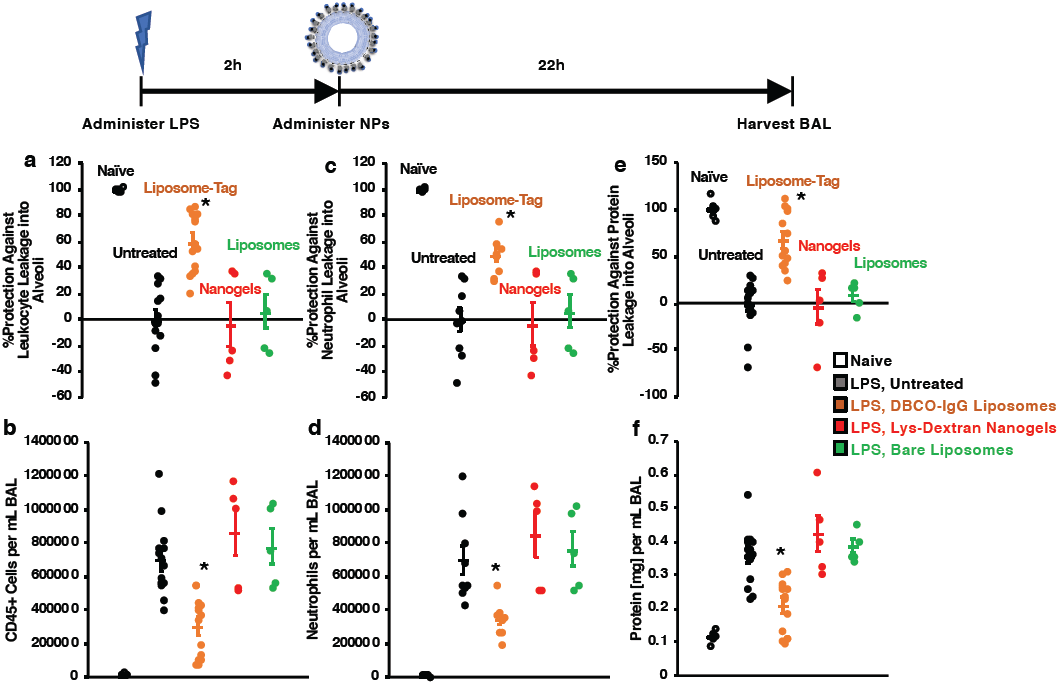
Therapeutic Effects of NAPs in Model Acute Respiratory Distress. Timeline: Nanoparticles or vehicle were administered intravenously two hours after nebulized LPS administration. Bronchoalveolar lavage (BAL) fluid was harvested 22 hours after liposome or vehicle administration. (a-b) Concentration of leukocytes (CD45-positive cells) in BAL fluid, represented as protection against infiltration into BAL (a) or absolute concentration (b). (c-d) Concentration of neutrophils (Ly6G-positive cells) in BAL fluid, represented as protection against infiltration into BAL (c) or absolute concentration (d). (e-f) Concentration of protein in BAL fluid (reflecting quantity of lung edema), represented as protection against edema (e) or absolute concentration (f). (*\* = p*<.*01*).

DBCO(20X)-IgG liposomes, LDNGs, and bare liposomes were compared for effects on vascular permeability in model ARDS. NPs were administered as an IV bolus (20 mg per kg body weight) two hours after nebulized LPS administration. As in untreated mice, BAL fluid was harvested and analyzed at 24 hours after exposure to nebulized LPS. Bare liposomes or LDNGs did not have significant effects on vascular injury induced by nebulized LPS, as measured by either leukocyte or protein concentration in BAL fluid (Figure 6, red and green bars). DBCO(20X)-IgG liposomes, however, had a significant salient effect on both protein leakage and cellular infiltration in the BAL (Figure 6, brown bars). With DBCO(20X)-IgG liposomes administered two hours after nebulized LPS, CD45-positive cells and neutrophils in BAL were reduced to concentrations of 3.04×10^5^ and 3.48×10^5^ cells per mL, respectively. Protein concentration in the BAL was reduced to 0.21 mg/mL by DBCO(20X)-IgG liposome treatment. As measured by protection against cellular or protein leakage, relative to untreated mice, DBCO(20X)-IgG liposomes provided 59.6% protection against leukocyte leakage, 49.7% protection against neutrophil leakage, and 67.4% protection against protein leakage. DBCO(20X)-IgG liposomes, without any drug, altered the course of inflammatory lung injury to limit protein and leukocyte edema in the alveoli.

Our results with DBCO(20X)-IgG liposomes indicate that some NAPs can interfere with neutrophil extravasation into the alveoli and thus limit edema following inflammatory injury. However, our results with LDNGs show that tropism for marginated neutrophils is not alone sufficient to limit the neutrophil-mediated effects of inflammatory lung injury.

## Discussion

Neutrophils concentrate in the pulmonary vasculature during either systemic or pulmonary inflammation.^10,11,14,17,18^ These marginated neutrophils can recognize and engulf bacteria.^17,18,35^ Therefore, neutrophils surveil the vasculature for potentially pathogenic foreign species, with the pulmonary vasculature serving as a “surveillance hub” in the case of systemic or pulmonary infection and inflammation.^10,17,18,35^ Our results with *E. coli* are noteworthy in this context: When *E. coli* are stripped of functional properties by heat treatment, oxidation, and fixation, but maintain their structure, uptake of the bacteria in the lungs only occurs after systemically prompting neutrophils with an inflammatory signal, LPS. Inflammation thus leads to pulmonary uptake of the *E. coli*-shaped particles.

In large part, the overall outcome of this study is an accounting of nanoparticle structural properties that lead to recognition by “surveilling” neutrophils in the inflamed lungs, analogously to *E. coli* recognition by pulmonary neutrophils. Including different liposomal formulations, 23 nanoparticles were screened in our biodistribution studies comparing pulmonary nanoparticle uptake in naïve and LPS-inflamed mice. Thirteen different nanoparticles exhibited specificity for inflamed lungs over naïve lungs, with flow cytometry data indicating that at least three of those nanoparticle species specifically and avidly gather in neutrophils.

The thirteen nanoparticles with specificity for the inflamed lungs have a range of properties. Seven different proteins were used in the inflammation-specific particles. The particles have sizes ranging from ∼75 nm to ∼350 nm, include both spheres and rods, and have a range of zeta potentials. However, our analyses classify the inflammation-specific nanoparticles as; 1) nanoparticles with structure based on hydrophobic interactions between proteins; 2) nanoparticles with structure based on non-site-specific protein crosslinking; 3) nanoparticles based on charge interactions between proteins. Put broadly, these three classes can all be grouped as structures based on *protein agglutination*, without regard for site-specific interactions or symmetry in the resulting protein superstructure.

We define the term **n**anoparticles with **a**gglutinated **p**roteins (NAPs) to indicate that particles with tropism for pulmonary marginated neutrophils during inflammation share commonalities in supramolecular organization. We identify NAPs as a broad class, rather than a single particle type. Accordingly, we have presented diverse NAP designs, implying a diversity of potential NAP-based strategies for targeted treatment and diagnosis of ARDS and other inflammatory disorders in which marginated neutrophils play a role (*e.g.* local infections or thrombotic disorders).^10,12,13,17,18^ The diversity of NAPs will allow versatile options for engineering neutrophil-specific drug delivery strategies to accommodate different pathologies.

In contrast to NAPs, three particles (adenovirus, AAVs, and ferritin) characterized by highly symmetric arrangement of protein subunits into a protein superstructure^44-46^ did not accumulate in the inflamed neutrophil-rich lungs. These three particles have evolved structures that lead to prolonged circulation or evasion of innate immunity in mammals.^53-56^ It is conceivable that neutrophils more effectively recognize less patterned and more variable protein arrangements that may better parallel the wide variety of structures presented by the staggering diversity of microbes against which neutrophils defend.^20,35^

To support our conclusions regarding supramolecular organization and neutrophil tropism, we re-engineered liposomes, particles with no intrinsic neutrophil tropism, to behave like NAPs. Protein arrangement on the surface of DBCO-IgG liposomes was predicted to recapitulate protein agglutination seen in NAPs based on hydrophobic interactions. Introduction of DBCO to IgG entails conjugation of a highly hydrophobic moiety^57^ to hydrophilic residues on the IgG. Replacing DBCO with the less hydrophobic modifying group used in SATA-maleimide conjugation^58^ abrogates the inflammation specificity observed with DBCO-IgG liposomes. Likewise, titrating down the amount of DBCO on the IgG, thus limiting the hydrophobic groups on the protein, also ratchets down the targeting behavior of the DBCO-IgG liposomes. Our data therefore points towards hydrophobic interactions between proteins on the liposome surface being a determinant in liposome uptake in neutrophils in the inflamed lungs. Essentially, the DBCO-IgG liposomes may reproduce the hydrophobic interaction structural motif seen in NAPs produced by protein gelation (*i.e.* LDNGs).

NAP-liposomes may be particularly attractive for future clinical translatability.

Liposomes are prominent among FDA-approved nanoparticle drug carriers.^59^ Further, even without cargo drugs, NAP-liposomes conferred significant therapeutic effects in a mouse model of severe ARDS. LDNGs, despite high levels of uptake in inflamed lungs, did not have the same therapeutic effect as the NAP-liposomes. This result suggests that the composition of the liposomes may be important for their therapeutic effect.

Among possible mechanisms for the therapeutic effect, we note that lipid rafts are major signaling hubs in neutrophils.^60,61^ The lipid content of the NAP-liposomes (particularly the cholesterol content) may modulate neutrophil lipid rafts dependent on cholesterol.

We have also observed that neutrophil content in the inflamed alveoli is markedly reduced by NAP-liposomes. In this context, we note published work demonstrating that certain nanoparticles, in a still undetermined manner dependent on particle composition, can drive redistribution of neutrophils from the lungs to the liver.^52^

As a major corollary, our findings indicate many protein-based or protein-incorporating nanoparticles developed for therapeutic applications may accumulate in inflamed lungs, even when those nanoparticles were designed to accumulate elsewhere. The variety of protein nanostructures accumulating in inflamed lungs in our data includes particles that have been investigated as targeted drug delivery vehicles where marginated neutrophils are not the intended site of accumulation.^36,38,47,48,62^ The patterns in our data indicate that future studies may reveal additional nanoparticles that accumulate in the lungs following inflammatory insult. This study therefore serves as evidence that inflammatory challenges may prompt profound off-target changes in the biodistributions of nanomaterials, including dramatic shunting of nanoparticles and any associated drug payload to the lungs. The nanoparticle targeting profiles documented in naïve or, for instance, tumor model studies may be overturned by, for instance, bacterial infection in a patient receiving the nanoparticle.

In conclusion, supramolecular organization in nanoparticle structure predicts nanoparticle uptake in pulmonary marginated neutrophils during acute inflammation. Specifically, nanoparticles with agglutinated protein (NAPs) accumulate in marginated neutrophils, while nanoparticles with more symmetric protein organization do not. NAP tropism for neutrophils allowed us to develop NAPs as diagnostics and therapeutics for ARDS, and even to demonstrate NAP uptake in inflamed human lungs. Future work may more deeply explore therapeutic effects of NAPs in ARDS and other diseases in which neutrophils play key roles. This study also obviates future testing of supramolecular organization as a variable in *in vivo* behavior of nanoparticles, including screens of tropism for other pathologies and cell types. These studies could in turn guide engineering of new particles with intrinsic cell tropisms, as with our engineering of NAP-liposomes with neutrophil tropism. These “targeting” behaviors, requiring no affinity moieties, may apply to a wide variety of nanomaterials. But our current findings with neutrophil-tropic NAPs indicate that many protein-based and protein-coated nanoparticles could be untapped resources for treatment and diagnosis of devastating inflammatory disorders like ARDS.

## Methods

### Lysozyme-Dextran Nanogel Synthesis

Lysozyme-dextran nanogels (LDNGs) were synthesized as previously described.^37,42^ 70 kDa rhodamine-dextran or FITC-dextran (Sigma) and lysozyme from hen egg white (Sigma) were dissolved in deionized and filtered water at a 1:1 or 2:1 mol:mol ratio, and pH was adjusted to 7.1 before lyophilizing the solution. For Maillard reaction between lysozyme and dextran, the lyophilized product was heated for 18 hours at 60°C, with 80% humidity maintained via saturated KBr solution in the heating vessel. Dextran-lysozyme conjugates were dissolved in deionized and filtered water to a concentration of 5 mg/mL, and pH was adjusted to 10.70 or 11.35. Solutions were stirred at 80°C for 30 minutes. Diameter of LDNGs was evaluated with dynamic light scattering (DLS, Malvern) after heat gelation. Particle suspensions were stored at 4°C.

### Crosslinked Protein Nanoparticle Synthesis

Crosslinked protein nanoparticles and nanorods were prepared using previously reported electrohydrodynamic jetting techniques.^63^ The protein nanoparticles were prepared using bovine serum albumin, human serum albumin, human lysozyme, human transferrin, or human hemoglobin (all proteins were purchased from Sigma). Protein nanorods were prepared using chemically modified human serum albumin.

For electrohydrodynamic jetting, protein solutions were prepared by dissolving the protein of interest at a 7.5 w/v% (or 2.5 w/v% for protein nanorods) concentration in a solvent mixture of DI water and ethylene glycol with 4:1 (v/v) ratio. The homo-bifunctional amine-reactive crosslinker, O,O′-bis[2-(N-succinimidyl-succinylamino)ethyl]polyethylene glycol with molecular weight of 2kDa (NHS-PEG-NHS, Sigma) was mixed with the protein solution at 10 w/w%. Protein nanoparticles were kept at 37°C for 7 days for completion of the crosslinking reaction. The as-prepared protein nanoparticles were collected in PBS buffer and their size distribution was analyzed using dynamic light scatting (DLS, Malvern).

### Glutamate-Tagged Green Fluorescent Protein Nanoparticle Synthesis

Glutamic acid residues (E20-tag) were inserted at the C-terminus of enhanced green fluorescent protein (EGFP) through restriction cloning and site-directed mutagenesis as previously reported.^64^ Proteins were expressed in an *E. coli* BL21 strain using standard protein expression protocol. Briefly, protein expression was carried out in 2xYT media with an induction condition of 1 mM IPTG and 18 °C for 16 h. At this point, the cells were harvested, and the pellets were lysed using 1% Triton-X-100 (30 min, 37 °C)/DNase-I treatment (10 minutes). Proteins were purified using HisPur cobalt columns. After elution, proteins were preserved in PBS buffer. The purity of native proteins was determined using 8% SDS-PAGE gel.

Polymers (PONI) were synthesized by ring-opening metathesis polymerization using third generation Grubbs’ catalyst as previously described.^65^ In brief, solutions in dichloromethane of guanidium functionalized monomer and Grubbs’ catalyst were placed under freeze thawing cycles for degassing. After warming the solutions to room temperature, the degassed monomer solutions were administrated to degassed catalyst solutions and allowed to stir for 30 minutes. The polymerization reaction was terminated by the addition of excess ethyl vinyl ether. The reaction mixture was further stirred for another 30 min. The resultant polymers were precipitated from excess hexane or diethyl ether anhydrous, filtered, washed and dried under vacuum to yield a light-yellow powder. polymers were characterized by ^1^H NMR and gel permeation chromatography (GPC) to assess chemical compositions and molecular weight distributions, respectively. Subsequent to deprotection of Boc functionalities, polymer was dissolved in the DCM with the addition of TFA at 1:1 ratio. The reaction was allowed to stir for 4 hours and dried under vacuum. Excess TFA was removed by azeotropic distillation with methanol. Afterwards, the resultant polymers were re-dissolved in DCM and precipitated in anhydrous diethyl ether, filtered, washed and dried. Polymers were then dissolved in water and transferred to Biotech CE dialysis tubing membranes with a 3000 g/mol cutoff and dialyzed against RO water (2-3 days). The polymers were then lyophilized dried to yield a light white powder.

PONI polymer/E-tag protein nanocomposites (PPNCs) were prepared in polypropylene microcentrifuge tubes (Fisher) through a simple mixing procedure. 0.5625 nmol of 54 kDa PONI was incubated with 0.45 nmol of EGFP at room temperature for 10 minutes prior to dilution to 200 µL in sterile PBS and subsequent injection. Similarly, 0.9 nmol of Arginine-tagged gold nanoparticles, prepared as described,^43^ were combined with 0.45 nmol of EGFP to prepare EGFP/gold nanoparticle complexes.

### Liposome Preparation

Azide-functionalized liposomes were prepared by thin film hydration techniques, as previously described.^47^ The lipid film was composed of 58 mol% DPPC (1,2-dipalmitoyl-sn-glycero-3-phosphocholine), 40 mol% cholesterol, and 2 mol% azide-PEG_2000_-DSPE (all lipids from Avanti). 0.5 mol% Top Fluor PC (1-palmitoyl-2-(dipyrrometheneboron difluoride) undecanoyl-sn-glycero-3-phosphocholine) was added to prepare fluorescent liposomes. 0.2 mol% DTPA-PE (1,2-distearoyl-sn-glycero-3-phosphoethanolamine-N-diethylenetriaminepentaacetic acid) was added to prepare liposomes with capacity for radiolabeling with ^111^In. Lipid solutions in chloroform, at a total lipid concentration of 20 mM, were dried under nitrogen gas, then lyophilized for 2 hours to remove residual solvent. Dried lipid films were hydrated with Dulbecco’s phosphate buffered saline (PBS). Lipid suspensions were passed through 3 freeze– thaw cycles using liquid N_2_/50°C water bath then extruded through 200 nm cutoff track-etched polycarbonate filters in 10 cycles. DLS assessed particle size after extrusion and after each subsequent particle modification. Liposome concentration following extrusion was assessed with Nanosight nanoparticle tracking analysis (Malvern).

For conjugation to liposomes, rat IgG was modified with dibenzylcyclooctyne-PEG_4_-NHS ester (DBCO, Jena Bioscience). IgG solutions (PBS) were adjusted to pH 8.3 with 1 M NaHCO_3_ buffer and reacted with DBCO for 1 hour at room temperature at molar ratios of 2.5:1, 5:1, 10:1, or 20:1 DBCO:IgG. Unreacted DBCO was removed after reaction via centrifugal filtration against 10 kDa cutoff filters (Amicon). Efficiency of DBCO-IgG reaction was assessed optically, with absorbance at 280nm indicating IgG concentration and absorbance at 309nm indicating DBCO concentration. Spectral overlap of DBCO and IgG absorbance was noted by correcting absorbance at 280nm according to *Abs*_280C_ = *Abs*_280_ − 1.089 × *Abs*_001_. Molar IgG concentration was determined according to 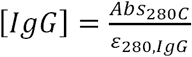, where *ε*_280,*IgG*_ is the IgG extinction coefficient at 280 nm, 204,000 L mol^-1^cm^-1^. Molar DBCO concentration was determined according to 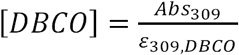, where *ε*_001,*DB*C*O*_ is the DBCO extinction coefficient at 309 nm, 12,000 L mol^-1^cm^-1^. Number of DBCO per IgG was determined as the ratio 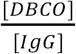. DBCO-modified IgG was incubated with azide liposomes at 200 IgG per liposome overnight at room temperature. Unreacted antibody was removed via size exclusion chromatography, and purified liposomes were concentrated to original volume against centrifugal filters (Amicon).

Maleimide liposomes were also prepared via lipid film hydration.^66^ Lipid films comprised 54% DPPC, 40% cholesterol, and 6% MPB-PE (1,2-dioleoyl-sn-glycero-3-phosphoethanolamine-N-[4-(p-maleimidophenyl) butyramide]), with lipids prepared, dried, resuspended, and extruded as described above for azide liposomes.

IgG was prepared for conjugation to maleimide liposomes by one-hour reaction of 10 SATA (N-succinimidyl S-acetylthioacetate) per IgG at room temperature in 0.5 mM EDTA in PBS. Unreacted SATA was removed from IgG by passage through 7 kDa cutoff gel filtration columns. SATA-conjugated IgG was deprotected by one-hour room temperature incubation in 0.05 M hydroxylamine in 2.5 mM EDTA in PBS. Excess hydroxylamine was removed and buffer was exchanged for 0.5 mM EDTA in PBS via 7 kDa cutoff gel filtration column. SATA-conjugated and deprotected IgG was added to liposomes at 200 IgG per liposomes for overnight reaction at 4°C. Excess IgG was removed by size exclusion column purification, as above for azide liposomes.

### Polystyrene Nanoparticle Preparation

150 nm carboxylate nanoparticles (Phosphorex) were exchanged into 50 mM MES buffer at pH 5.2 via gel filtration column. N-Hydroxysulfosuccinimide (sulfo-NHS) was added to the particles at 0.275 mg/mL, prior to incubation for 3 minutes at room temperature. EDCI was then added to the particles at 0.1 mg/mL, prior to incubation for 15 minutes at room temperature. IgG was added to the particle mixture at 200 IgG per nanoparticle, prior to incubation for 3 hours at room temperature while vortexing. For radiotracing, ^125^I-labeled IgG was added to the reaction at 5% of total IgG mass. The IgG/particle mixture was diluted with 10-fold volume excess of pH 5.2 MES buffer and the diluted mixture was centrifuged at 12000xg for 3 minutes. Supernatant was discarded and PBS with 0.05% BSA was added at desired volume before resuspending the particles via sonication probe sonication (three pulses, 30% amplitude). Particle size was assessed via DLS after resuspension, and particles were used immediately after DLS assessment.

### E. Coli Preparation

TOP10 *E. coli* were grown overnight in Terrific broth with ampicillin. Bacteria were heat-inactivated by 20-minute incubation at 60°C, then fixed by overnight incubation in 4% paraformaldehyde. After fixation, bacteria were pelleted by centrifugation at 1000xg for 10 minutes. Pelleted bacteria were washed three times in PBS, prior to resuspension by pipetting. Bacterial concentration was verified by optical density at 600 nm, prior to radiolabeling as described for nanoparticles below. Bacteria were administered in mice (7.5×10^7^ colony forming units in a 100 µL suspension per mouse).

### Protein, Nanoparticle, and Bacteria Iodination

Protein, horse spleen ferritin nanocages (Sigma), or adeno-associated virus (empty capsids, serotype 8) were prepared in PBS at concentrations between 1 and 2 mg/mL in volumes between 100 and 200 µL. Films of oxidizing agent were prepared in borosilicate tubes by drying 300 µL of 0.5 mg/mL Iodogen (Perkin-Elmer, chloroform solution) under nitrogen gas. Alternatively, Iodobeads (Perkin-Elmer) were added to borosilicate tubes (one per reaction). Protein solutions were added to coated or bead-containing tubes, before addition of Na^125/131^I at 25 µCi per 100 µg of protein. Protein was incubated with radioiodine at room temperature for 5 minutes under parafilm in a ventilated hood. Iodide-protein reacottions were terminated by purifying protein solutions through a 7 kDa cutoff gel column (Zeba). Additional passages through gel filtration columns or against centrifugal filters (Amicon, 10 kDa cutoff) were employed to remove free iodine, assuring that >95% of radioactivity was associated with protein.

Lysozyme-dextran nanogels, crosslinked protein nanoparticles, *E. coli*, or adenovirus were similarly iodinated. At least 100 µL of particle suspension was added to a borosilicate tube containing two Iodobeads, prior to addition of 100 µCi of Na^125^I per 100 µL of suspension. Particles were incubated with radioiodine and Iodobeads for 30 minutes at room temperature, with gentle shaking every 10 minutes. To remove free iodine, particle suspensions were moved to a centrifuge tube, diluted in ∼1 mL of buffer and centrifuged to pellet the particles (16000xg/30 minutes for nanogels, 16000xg/30 minutes for crosslinked protein particles, 10000xg/30 minutes for adenovirus, and 1000xg/10 minutes for *E. coli*). Supernatant was removed and wash/centrifugation cycles were repeated to assure >95% of radioactivity was associated with particles. Particles were resuspended by probe sonication (three pulses, 30% amplitude) for nanogels or crosslinked protein nanoparticles or pipetting for adenovirus or *E. coli*.

### Nanoparticle Labeling with ^111^In

^111^In labeling of nanoparticles followed previously described methods, with adaptation for new particles.^47^ All radiolabeling chelation reactions were performed using metal free conditions to prevent contaminating metals from interfering with chelation of ^111^In by DTPA or DOTA. Metals were removed from buffers using Chelex 100 metal affinity resin (Biorad, Laboratories, Hercules CA).

Lysozyme-dextran nanogels were prepared for chelation to ^111^In by conjugation to S-2-(4-Isothiocyanatobenzyl)-1,4,7,10-tetraazacyclododecane tetraacetic acid (p-SCN-Bn-DOTA, Macrocyclics). Nanogels were moved to metal free pH 8.3 1 M NaHCO_3_ buffer by three-fold centrifugation (16000xg for 15 minutes) and pellet washing with metal free buffer. p-SCN-Bn-DOTA was added to nanogels at 1:25 mass:mass ratio, prior to reaction for 30 minutes at room temperature. Free p-SCN-Bn-DOTA was removed by three-fold centrifugal filtration against 10 kDa cutoff centrifugal filters, with resuspension of nanogels in metal-free pH 4 citrate buffer after each centrifugation.

DOTA-conjugated nanogels or DTPA-containing liposomes in pH 4 citrate buffer were combined with ^111^InCl_3_ for one-hour chelation at 37°C. Nanoparticle/^111^InCl_3_ mixtures were treated with free DTPA (1 mM final concentration) to remove ^111^In not incorporated in nanoparticles. Efficiency of ^111^In incorporation in nanoparticles was assessed by thin film chromatography (aluminum/silica strips, Sigma) with 10 µM EDTA mobile phase. Chromatography strips were divided between origin and mobile front and the two portions of the strip were analyzed in a gamma counter to assess nanoparticle-associated (origin) vs. free (mobile front) ^111^In. Free ^111^In was separated from nanoparticles by centrifugal filtration and nanoparticles were resuspended in PBS (liposomes) or saline (nanogels). For SPECT/CT imaging experiments (see *SPECT/CT Imaging* methods below) with nanogels, 80 µCi of ^111^In-labeled nanogels, used within one day ^111^In labeling as described above, were administered to each mouse. For tracing ^111^In-labeled liposomes in biodistribution studies, liposomes were labeled with 50 µCi ^111^In per µmol of lipid.

### Nanoparticle Tracing in Inflammatory Disease Mouse Models

Nanoparticle or protein biodistributions were tested by injecting radiolabeled nanoparticles or protein (suspended to 100 µL in PBS or 0.9% saline at a dose of 2.5 mg/kg with tracer quantities of radiolabeled material) in C57BL/6 male mice from Jackson Laboratories. Biodistributions in naïve mice were compared to biodistributions in several injury models. Biodistribution data were collected at 30 minutes after nanoparticle or protein injection, unless otherwise stated, as in pharmacokinetics studies. Briefly, blood was collected by vena cava draw and mice were sacrificed via terminal exsanguination and cervical dislocation. Organs were harvested and rinsed in saline, and blood and organs were examined for nanoparticle or protein retention in a gamma counter (Perkin-Elmer). Nanoparticle or protein retention in harvested organs was compared to measured radioactivity in injected doses. For calculations of nanoparticle or protein concentration in organs, quantity of retained radioactivity was normalized to organ weights.

Mice subject to intravenous LPS injury were anesthetized with 3% isoflurane before administration of LPS from *E. coli* strain B4 at 2 mg/kg in 100 µL PBS via retroorbital injection. After five hours, mice were anesthetized with ketamine-xylazine (10 mg/kg ketamine, 100 mg/kg xylazine, intramuscular administration) and administered radiolabeled nanoparticles or protein via jugular vein injection to determine biodistributions as described above. For mice subject to intratracheal (IT) LPS injury, B4 LPS was administered to mice (anesthetized with ketamine/xylazine) at 1 mg/kg in 50 µL of PBS via tracheal catheter, followed by 100 µL of air.^66^ Biodistributions of lysozyme-dextran nanogels in IT-LPS-injured mice were assessed as above 16 hours after LPS administration. Biodistributions of liposomes in IT-LPS-injured mice were assessed at 1, 2, or 6 hours after LPS administration. Mice subject to footpad LPS administration were provided B4 LPS at 1 mg/kg in 50 µL PBS via footpad injection. Biodistributions of lysozyme-dextran nanogels were obtained at 6 or 24 hours after footpad LPS administration.

Lysozyme-dextran nanogel biodistributions were also traced in a mouse model of cardiogenic pulmonary edema.^50^ To establish edema, mice were anesthetized with ketamine/xylazine and administered propranolol in saline (3 µg/mL) via jugular vein catheter at 83 µL/min over 120 minutes. Lysozyme-dextran nanogel biodistributions were subsequently assessed as above.

### Single Cell Suspension Flow Cytometry

Single cell suspensions were prepared from lungs for flow cytometric analysis of cell type composition of the lungs and/or nanoparticle distribution among different cell types in the lungs. C57BL/6 male mice were anesthetized with ketamine/xylazine (10 mg/kg ketamine, 100 mg/kg xylazine, intramuscular administration) prior to installation of tracheal catheter secured by suture. After sacrifice by terminal exsanguination via the vena cava, lungs were perfused by right ventricle injection of ∼10 mL of cold PBS. The lungs were then infused via the tracheal catheter with 1 mL of a digestive enzyme solution consisting of 5 U/mL dispase, 2.5 mg/mL collagenase type I, and 1 mg/mL of DNAse I in cold PBS. Immediately after infusion, the trachea was sutured shut while removing the tracheal catheter. The lungs with intact trachea were removed via thoracotomy and kept on ice prior to manual disaggregation. Disaggregated lung tissue was aspirated in 2 mL of digestive enzyme solution and incubated at 37°C for 45 minutes, with vortexing every 10 minutes. After addition of 1 mL of fetal calf serum, tissue suspensions were strained through 100 µm filters and centrifuged at 400xg for 5 minutes. After removal of supernatant, the pelleted material was resuspended in 10 mL of cold ACK lysing buffer. The resulting suspensions were strained through 40 µm filter and incubated for 10 minutes on ice. The suspensions were centrifuged at 400xg for 5 minutes and the resulting pellets were rinsed in 10 mL of FACS buffer (2% fetal calf serum and 1 mM EDTA in PBS). After centrifugation at 400xg for 5 minutes, the rinsed cell pellets were resuspended in 2% PFA in 1 mL FACS buffer for 10 minutes incubation. The fixed cell suspensions were centrifuged at 400xg for 5 minutes and resuspended in 1 mL of FACS buffer.

For analysis of intravascular leukocyte populations in naïve and inflamed lungs, mice received an intravenous injection of FITC-conjugated anti-CD45 antibody five minutes prior to sacrifice and preparation of single cell suspensions as described above. Populations of intravascular vs. extravascular leukocytes were assessed by subsequent stain of fixed cell suspensions with PerCP-conjugated anti-CD45 antibody and/or APC-conjugated clone 1A8 anti-Ly6G antibody. To accomplish staining of fixed cells, 100 µL aliquots of the cell suspensions described above were pelleted at 400xg for 5 minutes, then resuspended in labeled antibody diluted in FACS buffer (1:150 dilution for APC-conjugated anti-Ly6G antibody and 1:500 dilution for PerCP-conjugated anti-CD45 antibody). Samples were incubated with staining antibodies for 20 minutes at room temperature in the dark, diluted with 1 mL of FACS buffer, and pelleted at 400xg for 5 minutes. Stained pellets were resuspended in 200 µL of FACS buffer prior to immediate flow cytometric analysis on a BD Accuri flow cytometer. All flow cytometry data was gated to remove debris and exclude doublets. Control samples with no stain, obtained from naïve and IV-LPS-injured mice, established gates for negative/positive staining with FITC, PerCP, and APC. Single stain controls allowed automatic generation of compensation matrices in FCS Express software. Comparison of PerCP anti-CD45 signal with FITC anti-CD45 signal indicated intravascular vs. extravascular leukocytes. Comparison of APC anti-Ly6G signal with FITC anti-CD45 signal indicated intravascular vs. extravascular neutrophils, with PerCP and APC co-staining verifying identification of cells as neutrophils.

Similar staining and analysis protocols enabled identification of nanoparticle distribution among different cell types in the lungs. To enable fluorescent tracing, lysozyme-dextran nanogels contained FITC-dextran, DBCO-IgG liposomes contained green fluorescent Top Fluor PC lipid, and crosslinked albumin nanoparticles were labeled with NHS ester Alexa Fluor 488. Alexa Fluor 488 labeling of albumin nanoparticles was accomplished by incubation of the NHS ester fluorophore with nanoparticles at 1:25 mass:mass fluorophore:nanoparticle ratio for two hours on ice. Excess fluorophore was removed from nanoparticles by 3-fold centrifugation at 16000xg for 15 minutes followed by washing with PBS. Nanoparticles were administered at 2.5 mg/kg via jugular vein injection and circulated for 30 minutes, prior to preparation of single cell suspensions from lungs as above. Fixed single cell suspensions were stained with APC-conjugated anti-Ly6G or PerCP-conjugated anti-CD45 as above. Additional suspensions were stained with 1:150 dilution of APC-conjugated anti-CD31, in lieu of anti-Ly6G, to identify endothelial cells. Association of nanoparticles with cell types was identified by coincidence of green fluorescent signal with anti-CD45, anti-Ly6G, or anti-CD31 signal.

### SPECT/CT Imaging

As described previously,^47^ thirty minutes after injection of 80 µCi of ^111^In-labeled nanogels, anesthetized mice were sacrificed by cervical dislocation. Mice were placed into a MiLabs U-SPECT (Utrecht, Netherlands) scanner bed. A region covering the entire body was scanned for 90 min using listmode acquisition. The animal was then moved, while maintaining position, to a MiLabs U-CT (Utrecht, Netherlands) for a full-body CT scan using default acquisition parameters (240 µA, 50 kVp, 75 ms exposure, 0.75° step with 480 projections). For naïve mice and mice imaged after cardiogenic pulmonary edema, CT data was acquired as above without SPECT data. The SPECT data was reconstructed using reconstruction software provided by the manufacturer, with 400 µm voxels. The CT data were reconstructed using reconstruction software provided by the manufacturer, with 100 µm voxels. SPECT and CT data, in NIFTI format, were opened with ImageJ software (FIJI package). Background signal was removed from SPECT images by thresholding limits determined by applying Renyi entropic filtering, as implemented in ImageJ, to a SPECT image slice containing NG-associated ^111^In in the liver. Background-subtracted pseudo-color SPECT images were overlayed on CT images and axial slices depicting lungs were selected for display, with CT thresholding set to emphasize negative contrast in the airspace of the lungs.

ImageJ’s built-in 3D modeling plugin was used to co-register background-subtracted pseudo-color SPECT images with CT images in three-dimensional reconstructions. CT image thresholding was set in the 3D modeling tool to depict skeletal structure alongside SPECT signal. For three-dimensional reconstructions of lung CT images, thresholding was set, as above, for contrast emphasizing the airspace of the lungs, with thresholding values standardized between different CT images (*i.e.* identical values were used for naïve and edematous lungs). Images were cropped in a cylinder to exclude the airspace outside of the animal, then contrast was inverted, allowing airspace to register bright CT signal and denser tissue to register as dark background. Three-dimensional reconstructions of the lung CT data, and co-registrations of SPECT data with lung CT data, were generated as above with ImageJ’s 3D plugin applied to CT data cropped and partitioned for lung contrast. Quantification of CT attenuation employed ImageJ’s measurement tool iteratively over axial slices, with measurement fields of view manually set to contain lungs and exclude surrounding tissue.

### Therapeutic Efficacy of Nanoparticles in Nebulized LPS Model

Mice were exposed to nebulized LPS in a ‘whole-body’ exposure chamber, with separate compartments for each mouse (MPC-3 AERO; Braintree Scientific). To maintain adequate hydration, mice were injected with 1mL of sterile saline warmed to 37°C, intraperitoneally, immediately before exposure to LPS. LPS (L2630-100mg, Sigma Aldrich) was reconstituted in PBS to 10 mg/mL and stored at −80°C until use. Immediately before nebulization, LPS was thawed and diluted to 5 mg/mL with PBS. LPS was aerosolized via a jet nebulizer connected to the exposure chamber (NEB-MED H, Braintree Scientific, Inc.). 5 mL of 5 mg/mL LPS was used induce the injury. Nebulization was performed until all liquid was nebulized (∼20 minutes).

Liposomes or saline sham were administered via retro-orbital injections of 100 µL of suspension (25 mg/kg liposome dose) at 2 hours after LPS exposure. Mice were anesthetized with 3% isoflurane to facilitate injections. Bronchoalveolar lavage (BAL) fluid was collected 24 hours after LPS exposure, as previously described.^66^ Briefly, mice were anesthetized with ketamine-xylazine (10 mg/kg ketamine, 100 mg/kg xylazine, intramuscular administration). The trachea was isolated and a tracheostomy was performed with a 22-gauge catheter. The mice were euthanized via exsanguination. 0.8 mL of cold BAL buffer (0.5 mM EDTA in PBS) was injected into the lungs over ∼1min via the tracheostomy and then aspirated from the lungs over ∼1min. Injections/aspirations were performed three times for a total of 2.4mL of fluid added to the lungs. Recovery BAL fluid typically amounted to ∼2.0mL.

BAL samples were centrifuged at 300xg for 4 minutes. The supernatant was collected and stored at −80°C for further analysis. Protein concentration was measured using Bio-Rad DC Protein Assay, per manufacturer’s instructions. The cell pellet was fixed for flow cytometry as follows. 333 µL of 1.6% PFA in PBS was added to each sample. Samples were incubated in the dark at room temperature for 10 minutes, then 1 mL of BAL buffer was added. Samples were centrifuged at 400xg for 3min, the supernatant was aspirated, and 1 mL of FACS buffer (2% fetal calf serum and 1 mM EDTA in PBS) was added. At this point, samples were stored at 4°C for up to 1 week prior to flow cytometry analysis.

To stain for flow cytometry, samples were centrifuged at 300xg for 4 min, the supernatant was aspirated, and 100 µL of staining buffer was added. Staining buffer used was a 1:1000 dilution of stock antibody solution (APC anti-mouse CD45; Alexa Fluor 488 anti-mouse Ly6G, Biolegend) into FACS buffer. Samples were incubated with staining antibody for 30 minutes at room temperature in the dark. To terminate staining, 1 mL of FACS buffer was added, samples were centrifuged at 300xg for 4 minutes, and supernatant was aspirated. Cells were resuspended in 900 µL of FACS buffer and immediately analyzed via flow cytometry.

Flow cytometric analysis was completed with a BD Accuri flow cytometer as follows: Sample volume was set to 100 µL and flow rate was set to ‘fast’. Unstained and single-stained controls were used to set gates. Forward scatter (pulse area) vs. side scatter (pulse area) plots were used to gate out non-cellular debris. Forward scatter (pulse area) vs. forward scatter (pulse height) plots were used to gate out doublets. The appropriate fluorescent channels were used to determine stained vs. unstained cells. The gates were placed using unstained control samples. Single-stain controls were tested and showed there was no overlap/bleed-through between the fluorophores. Final analysis indicated the quantity of leukocytes (CD45-positive cells) and neutrophils (Ly6G-positive cells) in BAL samples.

### Nanoparticle Administration in Human Lungs

Human lungs were obtained after organ harvest from transplant donors whose lungs were in advance deemed unsuitable for transplantation. The lungs were harvested by the organ procurement team and kept at 4°C until the experiment, which was done within 24 hours of organ harvest. The lungs were inflated with low pressure oxygen and oxygen flow was maintained at 0.8 L/min to maintain gentle inflation.

Pulmonary artery subsegmental branches were endovascularly cannulated, then tested for retrograde flow by perfusing for 5 minutes with Steen solution containing a small amount of green tissue dye at 25 cm H_2_O pressure. The pulmonary veins through which efflux of perfusate emerged were noted, allowing collection of solutions after passage through the lungs. A 2 mL mixture of ^125^I-labeled lysozyme-dextran nanogels and ^131^I-labeled ferritin nanocages were injected through the arterial catheter. ∼100mL of 3% BSA in PBS was passed through the same catheter to rinse unbound nanoparticles. A solution of green tissue dye was subsequently injected through the same catheter. The cannulated lung lobe was dissected into ∼1 g segments, which were evaluated for density of tissue dye staining. Segments were weighed, divided into ‘high’, ‘medium’, ‘low’, and ‘null’ levels of dye staining, and measured for ^131^I and ^125^I signal in a gamma counter.

For experiments with cell suspensions derived from human lungs (chosen for research use according to the above standards), single cell suspensions were generously provided by the laboratory of Edward E. Morrisey at the University of Pennsylvania. Aliquots of 600,000 cells were pelleted at 400xg for 5 minutes and resuspended in 100 µL PBS containing different quantities of lysozyme-dextran nanogels synthesized with FITC-labeled dextran. Cells and nanogels were incubated at room temperature for 30 minutes before two-fold pelleting at 400xg with 1 mL PBS washes. Cells were re-suspended in 200 µL FACS buffer for staining with APC-conjugated anti-human CD45, applied by 20-minute incubation with a 1:500 dilution of the antibody stock. Cells were pelleted at 400xg for 5 minutes and resuspended in 200 µL PBS for immediate analysis with flow cytometry (BD Accuri). Negative/positive nanogel or anti-CD45 signal was established by comparison to unstained cells. Single-stained controls indicated no spectral overlap between FITC-nanogel fluorescence and anti-CD45 APC fluorescence.

### Circular Dichroism Spectroscopy

Proteins were prepared in deionized and filtered water at concentrations of 0.155 mg/mL for human albumin, 0.2 mg/mL for hen lysozyme, and 0.48 mg/mL for IgG. Crosslinked albumin nanoparticles, lysozyme-dextran nanogels, and IgG-coated liposome suspensions were prepared such that albumin, lysozyme, and IgG concentrations in the suspensions matched the concentrations of the corresponding protein solutions. Protein and nanoparticle solutions were analyzed in quartz cuvettes with 10 mm path length in an Aviv circular dichroism spectrometer. The instrument was equilibrated in nitrogen at 25°C for 30 minutes prior to use and samples were analyzed with sweeps between 185 and 285 nm in 1 nm increments. Each data point was obtained after a 0.333 s settling time, with a 2 s averaging time. CDNN^40^ software deconvolved CD data (expressed in millidegrees) via neural network algorithm assessing alignment of spectra with library-determined spectra for helices, antiparallel sheets, parallel sheets, beta turns, and random coil.

### 8-anilino-1-naphthalenesulfonic Acid Nanoparticle Staining

8-anilino-1-naphthalenesulfonic acid (ANSA) at 0.06 mg/mL was mixed with lysozyme, human albumin, or IgG at 1.5 mg/mL in PBS. For nanoparticle analysis, nanoparticle solutions were prepared such that albumin, lysozyme, and IgG concentrations in the suspensions matched the 1.5 mg/mL concentration of the protein solutions. Protein or nanoparticles and ANSA were reacted at room temperature for 30 minutes. Excess ANSA was removed from solutions by 3 centrifugations against 3 kDa cutoff centrifugal filters (Amicon). After resuspension to original volume, ANSA-stained protein/nanoparticle solutions/suspension were examined for fluorescence (excitation 375 nm, emission 400-600 nm) and absorbance (240-540 nm) maxima corresponding to ANSA.

### Histology

For imaging neutrophil content in naïve and IV-LPS-injured lungs, mice were intravenously injected with rat anti-mouse anti-Ly6G antibody (clone 1A8) and sacrificed 30 minutes later. Lungs were embedded in M1 medium, flash frozen, and sectioned in 10 µm slices. Sections were stained with PerCP-conjugated anti-rat secondary antibody and neutrophil-associated fluorescence was observed with epifluorescence microscopy. Similar procedures enabled histological imaging of lysozyme-dextran nanogels in IV-LPS-injured lungs. Nanogels synthesized with rhodamine-dextran were administered intravenously in injured mice 30 minutes prior to sacrifice. Lungs were sectioned as above and stained with clone 1A8 anti-Ly6G antibody, followed by Briliant Violet-conjugated anti-rat secondary antibody.

Sections of human lungs were obtained after ex vivo administration (see *Nanoparticle Administration in Human Lungs* above) of lysozyme-dextran nanogels synthesized with rhodamine dextran. Regions of tissue delineated as perfused and non-perfused, as determined by arterial administration of tissue dye as above, were harvested, embedded in M1 medium, flash frozen, and sectioned in 10 µm slices. Epifluorescence imaging indicated rhodamine fluorescence from nanogels, co-registered to autofluorescence indicating tissue architecture.

### Live Lung Imaging

A mouse was anesthetized with ketamine/xylazine five hours after intravenous administration of 2 mg/kg B4 LPS. A jugular vein catheter was fixed in place for injection of lysozyme-dextran nanogels, anti-CD45 antibody, and fluorescent dextran during imaging. In preparation for exposure of the lungs, a patch of skin on the back of the mouse, around the juncture between the ribcage and the diaphragm, was denuded.

While the mouse was maintained on mechanical ventilation, an incision at the juncture between the ribs and the diaphragm, towards the posterior, exposed a portion of the lungs. A coverslip affixed to a rubber o-ring was sealed to the incision by vacuum. The exposed portion of the mouse lung was placed in focus under the objective by locating autofluorescence signal in the “FITC” channel. With 100 ms exposure, fluorescent images from channels corresponding to violet, green, near red, and far red fluorescence were sequentially acquired. A mixture of rhodamine-dextran nanogels (2.5 mg/kg), Brilliant Violet-conjugated anti-CD45 antibody (0.8 mg/kg), and Alexa Fluor 647 labeled 70 kDa dextran (40 mg/kg) for vascular contrast was administered via jugular vein catheter and images were recorded for 30 minutes. Images were recorded in SlideBook software and opened in ImageJ (FIJI distribution) for composition in movies with co-registration of the four fluorescent channels.

### Animal and human study protocols

All animal studies were carried out in strict accordance with Guide for the Care and Use of Laboratory Animals as adopted by National Institute of Health and approved by University of Pennsylvania Institutional Animal Care and Use Committee (IACUC). Male C57BL/6J mice, 6-8 weeks old, were purchased from Jackson Laboratories. Mice were maintained at 22–26°C and on a 12/12 hour dark/light cycle with food and water ad libitum.

Ex vivo human lungs were donated from an organ procurement agency, Gift of Life, after determination the lungs were not suitable for transplantation into a recipient, and therefore would have been discarded if they were not used for our study. Gift of Life obtained the relevant permissions for research use of the discarded lungs, and in conjunction with the University of Pennsylvania’s Institutional Review Board ensured that all relevant ethical standards were met.

### Statistical Analysis

Error bars indicate standard error of the mean throughout. Significance was determined through paired t-test for comparison of two samples and ANOVA for group comparisons. Linear discriminant analysis and principal components analysis were completed in Gnu Octave scripts (adapted from https://www.bytefish.de/blog/pca_lda_with_gnu_octave/, and made available in full in the supplementary materials).

## Acknowledgments

This research was supported by NIH R01HL125462 (V.R.M.). The authors acknowledge funding from the Defense Threat Reduction Agency (HDTRA-1-15-0045, J.L., V.R.M.). V.M.R. acknowledges NIH EB022641. J.W.M. was supported by NIH T32HL07915 while conducting this research. O.A.M. received funding from the American Heart Association (19CDA345900001). Circular dichroism spectroscopy data were obtained at the University of Pennsylvania Biological Chemistry Resource Center. SPECT-CT images were obtained by Eric Blankenmyer in collaboration with the University of Pennsylvania Perelman School of Medicine Small Animal Imaging Facility. Single cell suspensions prepared from human lungs were generously provided by Edward E. Morrisey’s group at the University of Pennsylvania. The authors report no competing financial interests.

## Data Availability

Raw imaging, flow cytometry, gamma counter, and spectroscopy data supporting the findings of this study are available from the corresponding author upon reasonable request. All other data supporting the findings of this study are available within the paper and its supplementary information files.

## Author Contributions

J.W.M, P.N.P, N.H., L.R.W., D.C.L, Y.L., O.A.M., E.D.H., T.S., J.V.G., and J.N. designed and prepared nanoparticles used in the study. J.W.M., P.M.G., and R.Y.K. prepared *E. coli* for tracing in mice. J.W.M., P.N.P., L.R.W., O.A.M., P.M.G., I.J., J.N., and C.F.G. performed studies tracing nanoparticles in mice. J.W.M., P.N.P., M.H.Z., and M.E.Z. performed studies of therapeutic efficacy. J.W.M., P.N.P., and L.T.F. performed studies tracing nanoparticles in human lungs. J.W.M., L.T.F. and K.M.R. performed studies tracing nanoparticle uptake in isolated cells. J.W.M., P.N.P., V.R.M., and J.S.B. analyzed all data. The manuscript was written through contributions of all authors. All authors have given approval to the final version of the manuscript.

