## Supplemental Figures, Tables, and Methods for "Supramolecular Organization Predicts Protein Nanoparticle Delivery to Neutrophils for Acute Lung Inflammation Diagnosis and Treatment"

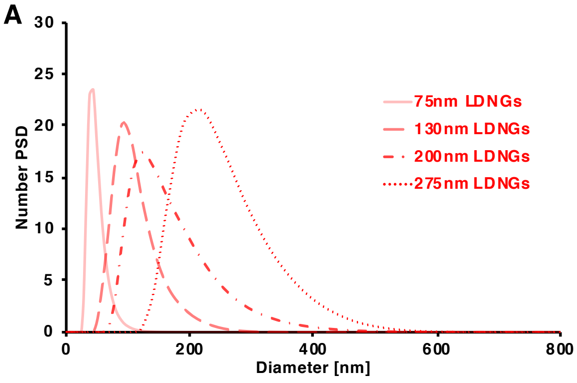


| Lysozyme-Dextran Nanogels | |
|---|---|
| Diameter | PDI |
| 73.21±1.28 nm | 0.181±0.053 |
| 136.38±3.60 nm | 0.100±0.022 |
| 199.44±1.81 nm | 0.111±0.011 |
| 274.50±6.44 nm | 0.155±0.062 |

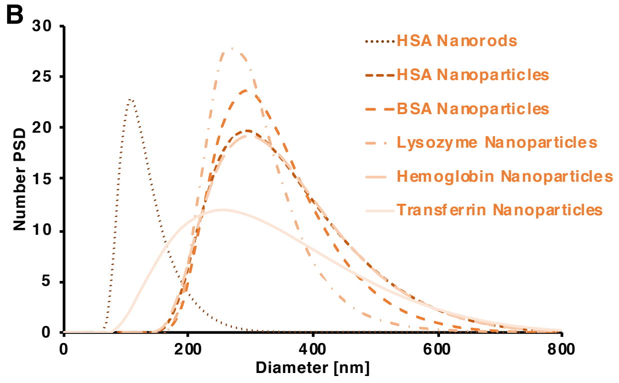


| Crosslinked Protein Nanoparticles | | |
|---|---|---|
| | Diameter | PDI |
| **HSA Nanorods** | 125.60±5.52 nm | 0.083±0.035 |
| **HSA NPs** | 317.82±3.60 nm | 0.144±0.052 |
| **BSA NPs** | 317.27±38.49 nm | 0.168±0.039 |
| **Lysozyme NPs** | 298.61±12.35 nm | 0.061±0.013 |
| **Hemoglobin NPs** | 328.08±16.08 nm | 0.080±0.010 |
| **Transferrin NPs** | 345.24±10.23 nm | 0.117±0.004 |

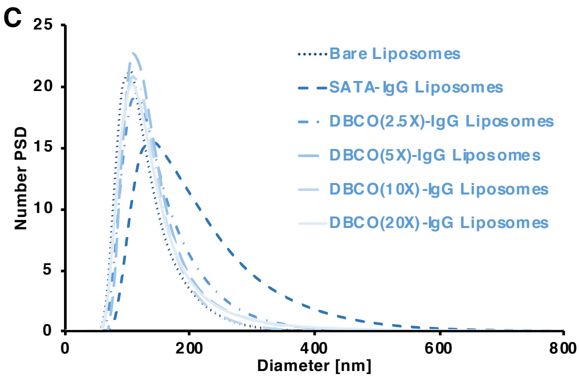


| Liposomes | | |
|---|---|---|
| | Diameter | PDI |
| **Bare Liposomes** | 103.63±8.66 nm | 0.091±0.007 |
| **SATA-IgG Liposomes** | 178.75±6.95 nm | 0.230±0.034 |
| **DBCO(2.5X)-IgG Lip.** | 136.60±5.66 nm | 0.158±0.032 |
| **DBCO(5X)-IgG Lip.** | 132.99±0.23 nm | 0.140±0.005 |
| **DBCO(10X)-IgG Lip.** | 122.93±7.56 nm | 0.085±0.013 |
| **DBCO(20X)-IgG Lip.** | 128.25±4.26 nm | 0.172±0.029 |

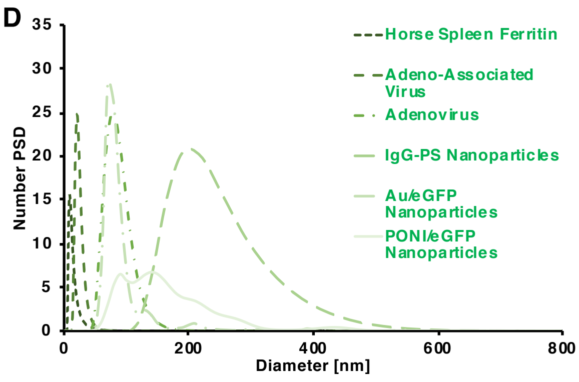


| Miscellaneous | | |
|---|---|---|
| | Diameter | PDI |
| **Horse Spleen Ferritin** | 12.04±0.35 nm | 0.054±0.002 |
| **Adeno-Associated Virus** | 23.22±0.98 nm | 0.014±0.001 |
| **Adenovirus** | 82.51±10.65 nm | 0.029±0.006 |
| **IgG-Polystyrene NPs** | 230.48±2.79 nm | 0.142±0.009 |
| **Au/eGFP NPs** | 88.95±1.56 nm | 0.136±0.036 |
| **PONI/eGFP NPs** | 158.93±6.16 nm | 0.173±0.025 |

### Supplementary Figure 1

*Dynamic Light Scattering or Nanosight Characterization of Tested Nanoparticles.* (a) Size measurements of lysozyme-dextran nanogel variants. Nanogels with 2:1 mol:mol rhodamine-dextran:lysozyme composition had a diameter of 73.21±1.28 nm, PDI 0.181±0.053. Nanogels with 1:1 mol:mol rhodamine-dextran:lysozyme composition had a diameter of 136.38±3.60 nm, PDI 0.100±0.022. Nanogels with 1:1 mol:mol FITC-dextran:lysozyme composition had a diameter of 199.44±1.81 nm, PDI 0.111±0.011. Nanogels with 1:1 mol:mol rhodamine-dextran:lysozyme composition, formed at pH 10.70, had a diameter of 274.50±6.44 nm, PDI 0.155±0.062. (b) Size measurements of crosslinked protein nanoparticle variants. Nanoparticles or nanorods were formed via co-jetting of PEG-NHS-ester-crosslinker with human albumin, bovine albumin, hen lysozyme, human hemoglobin, or human transferring. (c) Size measurements of variant liposome formulations. Bare liposomes had a diameter of 103.63±8.66 nm, PDI 0.091±0.007. Maleimide liposomes conjugated to SATA-functionalized IgG had a diameter of 176.75±6.95 nm, PDI 0.230±0.034. Azide-presenting liposomes conjugated to DBCO-functionalized IgG had diameters of approximately 130nm, with small variations registered for different DBCO densities on IgG. (d) Size measurements of other nanoparticles used in the study. Naturally occurring horse spleen ferritin, adeno-associated virus, and adenovirus diameters were confirmed by DLS. Horse spleen ferritin had a diameter of 12.04±0.35 nm, PDI 0.054±0.002. Adeno-associated virus had a diameter of 23.22±0.98 nm, PDI 0.014±0.001. Adenovirus had a diameter of 82.51±10.65 nm, PDI 0.029±0.006. Carboxylated polystyrene nanoparticles (initial diameter ~150nm) were conjugated to IgG via EDCI-mediated carboxy-amine reaction, yielding particles with diameter of 230.48±2.79 nm, PDI 0.142±0.009. Polyglutamate-tagged green fluorescent protein (eGFP) was combined with arginine-tagged gold nanoclusters (Au) or arginine-poly(oxanorborneneimide) (PONI), forming particles with diameter, as assessed by nanoparticle tracking analysis, of 88.95±1.56 nm (PDI 0.136±0.036) for Au-eGFP and 158.93±6.16 (PDI 0.173±0.025) for PONI-eGFP.

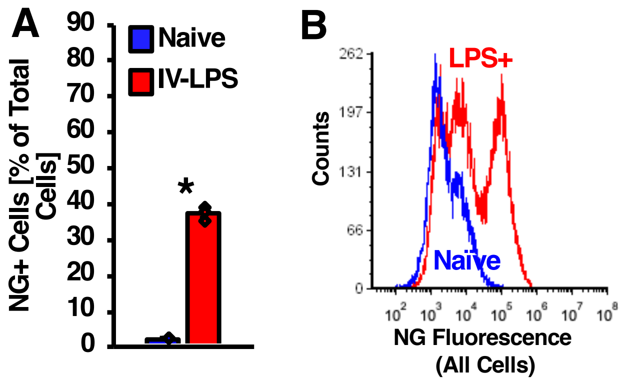


**Supplementary Figure 2**

*Flow Cytometric Characterization of Lysozyme-Dextran Nanogel Uptake in Naïve and Inflamed Lungs.* (a) Fluorescence from FITC-labeled lysozyme-dextran nanogels was measured in single cell suspensions prepared from mouse lungs harvested after 30 minutes nanogel circulation. With gates set as depicted in main text figure 2c, the number of cells positive for nanogel fluorescence increased between naïve and LPS-challenged lungs (** = p<.01)*. (b) Likewise, a population of high-fluorescence cells was detected in IV LPS-challenged lungs, but not naïve lungs.

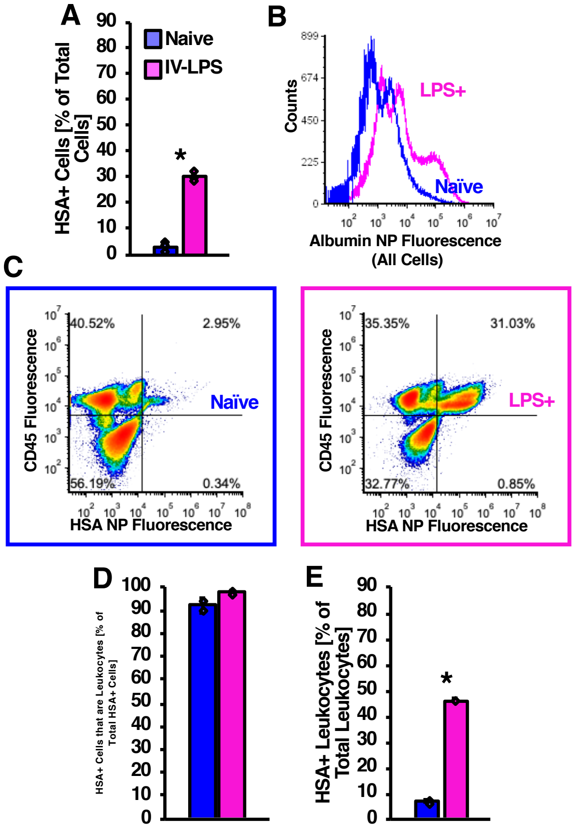


**Supplementary Figure 3**

*Flow Cytometric Characterization of Crosslinked Albumin Nanoparticle Uptake in Leukocytes in Naïve and Inflamed Lungs.* (a) Fluorescence from Alexa Fluor 498-labeled crosslinked albumin nanoparticles was measured in single cell suspensions prepared from mouse lungs harvested after 30 minutes nanoparticle circulation. With gates set as depicted in main text figure 2d, the number of cells positive for albumin nanoparticle fluorescence increased between naïve and LPS-challenged lungs. (b) A population of high-fluorescence cells was detected in IV LPS-challenged lungs, but not naïve lungs. (c) Fluorescence generated by CD45 staining, distinguishing leukocytes in single cell suspensions, plotted against human albumin nanoparticle fluorescence in single cell suspensions prepared from naïve and IV LPS-challenged lungs. (d) With gates set by the quadrants delineated in (c), correlation between nanoparticle fluorescence and CD45 staining indicated the percentage of albumin nanoparticle-bearing cells that were leukocytes as >90% in both naïve and IV LPS-challenged lungs. (e) Similar analysis indicated that the fraction of leukocytes containing albumin nanoparticles increased in LPS-challenged vs. naïve lungs. (** = p<.01*)

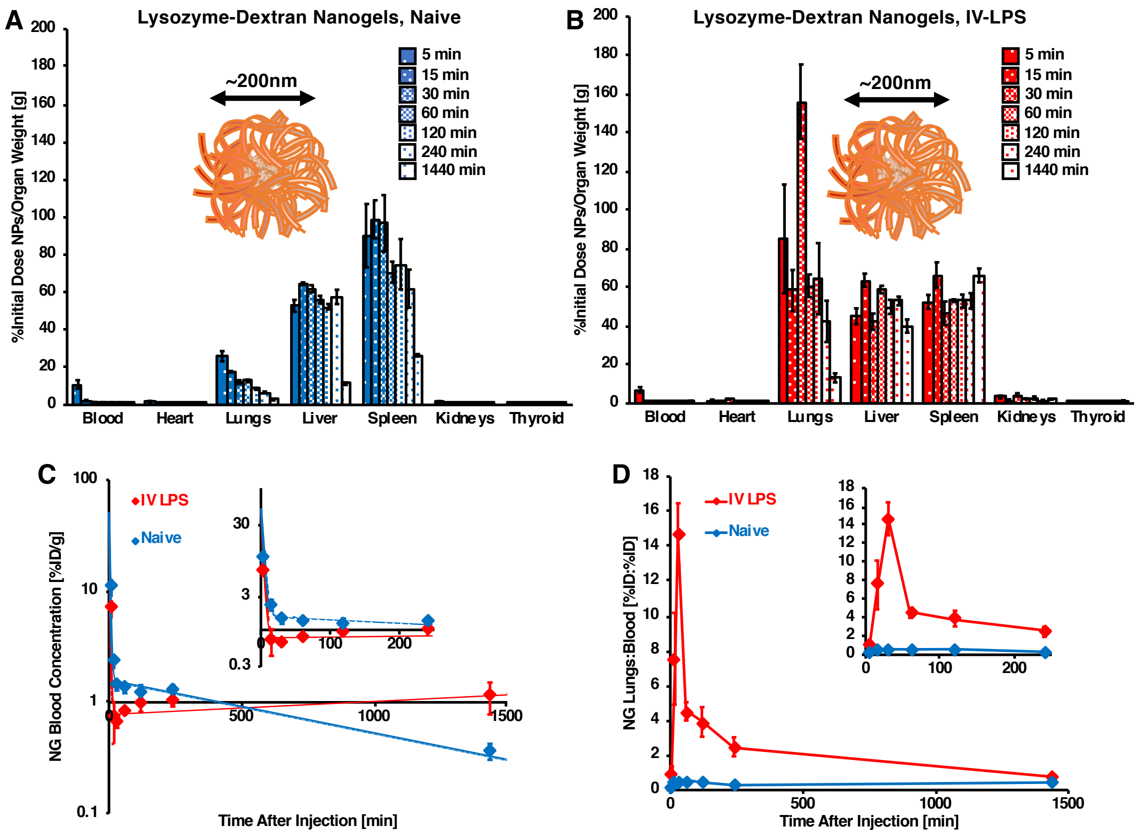


**Supplementary Figure 4**

*Pharmacokinetics of Lysozyme-Dextran Nanogels in Naïve and IV-LPS-Injured Mice.* Lysozyme-dextran nanogel biodistributions were determined at 5, 15, 30, 60, 120, 240, and 1440 minutes after bolus injection in naïve (a, blue) and IV LPS-challenged (b, red) mice. (c) Log-linear representation of nanogel clearance from the blood over 24 hours after injection (inset: 0-4 hour clearance data) indicating rapid clearance in both naïve (blue) and LPS-challenged (red) mice, with blood levels of nanogels being lower in LPS-challenged mice between 0 and 4 hours after injection, but greater in LPS-challenged mice at 24 hours after injection. (d) Nanogel lung uptake:blood level ratio in naïve (blue) and LPS-challenged (red) mice. Lungs:blood metric reaches a clear peak at 30 minutes after nanogel injection in LPS-injured mice (inset: 0-4 hour lungs:blood data).

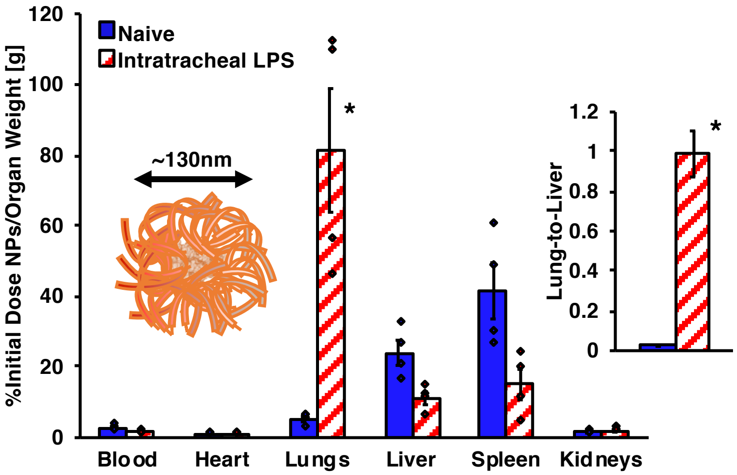


**Supplementary Figure 5**

*Biodistributions of Lysozyme-Dextran Nanogels in Naïve and Intratracheal LPS-Injured Mice.* As an alternative to intravenous LPS injection, mice were administered LPS via intratracheal (IT) instillation, prior to bolus dosing with lysozyme-dextran nanogels. As with IV LPS-injured mice, IT LPS injury led to dramatically increased pulmonary uptake of lysozyme-dextran nanogels, along with depression in hepatic and splenic uptake, relative to values in naïve mice (** = p<.001*).

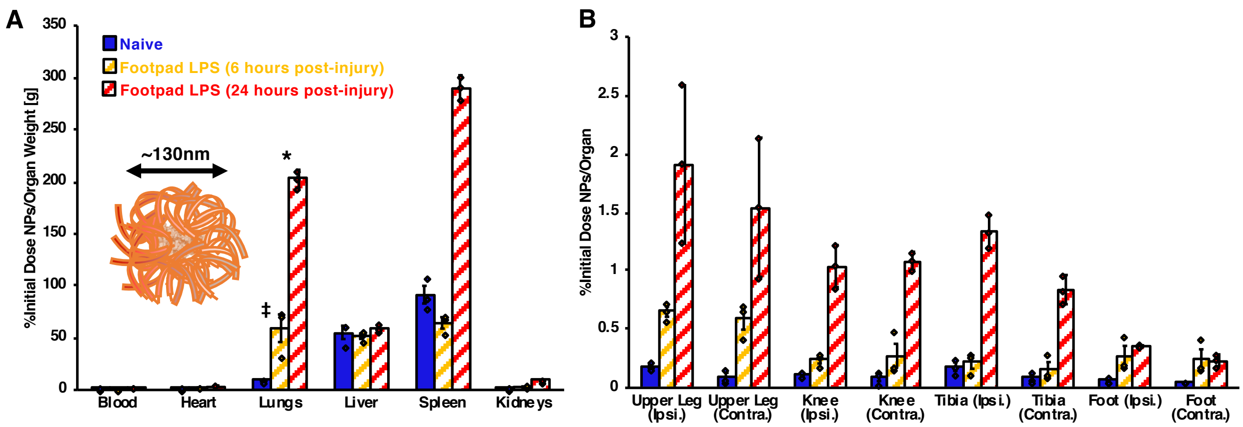


**Supplementary Figure 6**

*Biodistributions of Lysozyme-Dextran Nanogels After Footpad Administration of LPS.* As an additional mouse model for LPS-induced inflammation, LPS was dosed in the footpad, either 5 or 24 hours prior to dosing with lysozyme-dextran nanogels. Nanogel uptake in the lungs, legs, and feet increased with time after footpad LPS administration. Pulmonary uptake at both 5 and 24 hours after LPS was significantly increased relative to uptake in naïve mice (‡ = $p<.01$, * = $p<.001$). No significant differences were noted in nanogel uptake in ipsilateral vs. contralateral legs.

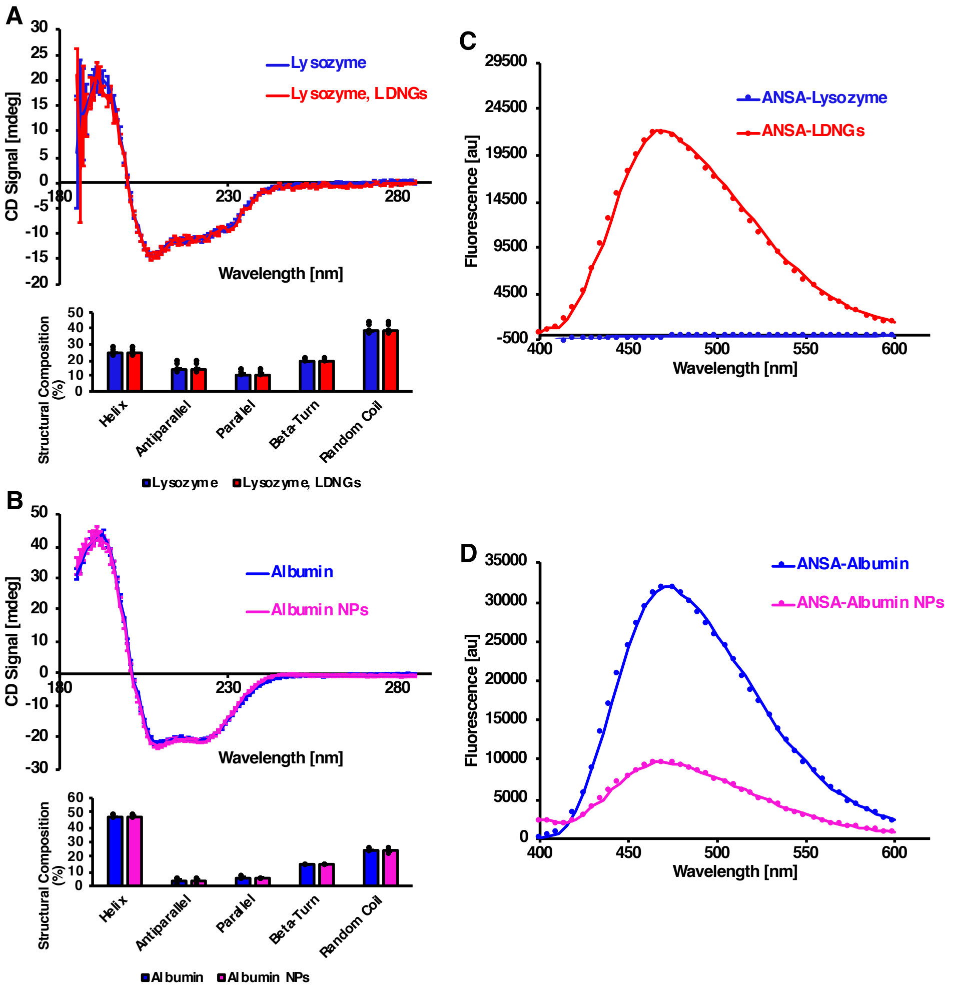


**Supplementary Figure 7**

*Circular Dichroism Spectroscopic Characterization of Protein Secondary Structure and ANSA Characterization of Hydrophobic Domain Accessibility for Lysozyme-Dextran Nanogels and Crosslinked Albumin Nanoparticles.* (a) Circular dichroism spectra for lysozyme-dextran nanogels and free lysozyme, with free lysozyme concentration set to match the concentration of lysozyme in the nanogels. Inset: neural network deconvolution of CD spectra indicating no differences in secondary structure composition between isolated lysozyme and lysozyme in nanogels. (b) Circular dichroism spectra for crosslinked human albumin nanoparticles and free human albumin, with free albumin concentration set to match the concentration of albumin in the nanoparticles. Inset: neural network deconvolution of CD spectra indicating no differences in secondary structure composition between isolated albumin and albumin in nanoparticles. (c) 8-anilino-1-naphthalenesulfonic acid (ANSA) staining of free lysozyme or lysozyme-dextran nanogels. Increased ANSA fluorescence indicates increased accessibility of hydrophobic domains in the nanogels, compared to lysozyme. (d) ANSA staining of free human albumin and albumin nanoparticles. Reduced ANSA fluorescence indicates lesser accessibility of hydrophobic domains in the nanoparticles, compared to albumin.

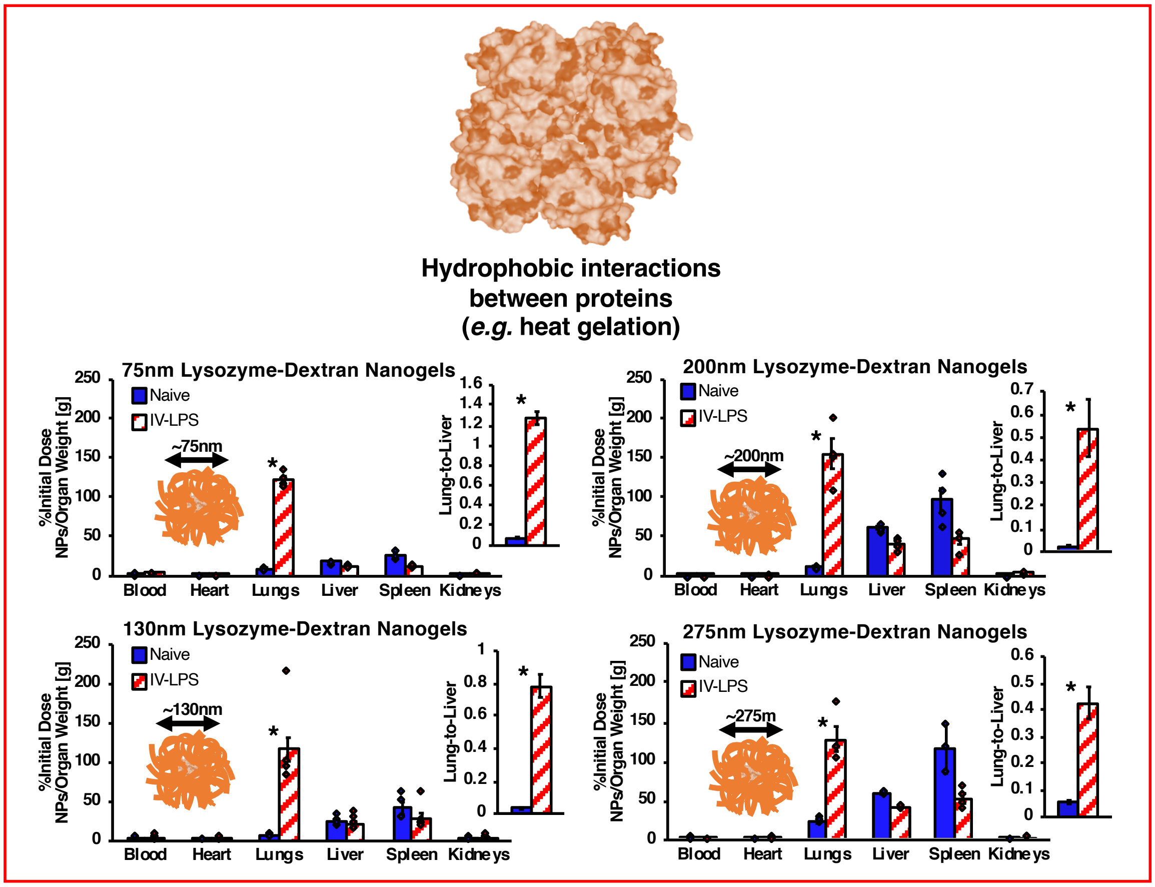


**Supplementary Figure 8**

*Biodistributions of Structural Variants of Lysozyme-Dextran Nanogels in Naïve and IV-LPS-Injured Mice.* Different lysozyme-dextran nanogel formulations, as described in supplementary figure 1a, were traced in naïve and intravenous LPS-challenged mice. LPS treatment enhanced pulmonary nanogel uptake for all nanogel variants. Lung, liver, and blood data for 75nm, 130nm, and 200nm nanogels are reproduced in figure 3a in the main text. (* = *p<.001*)

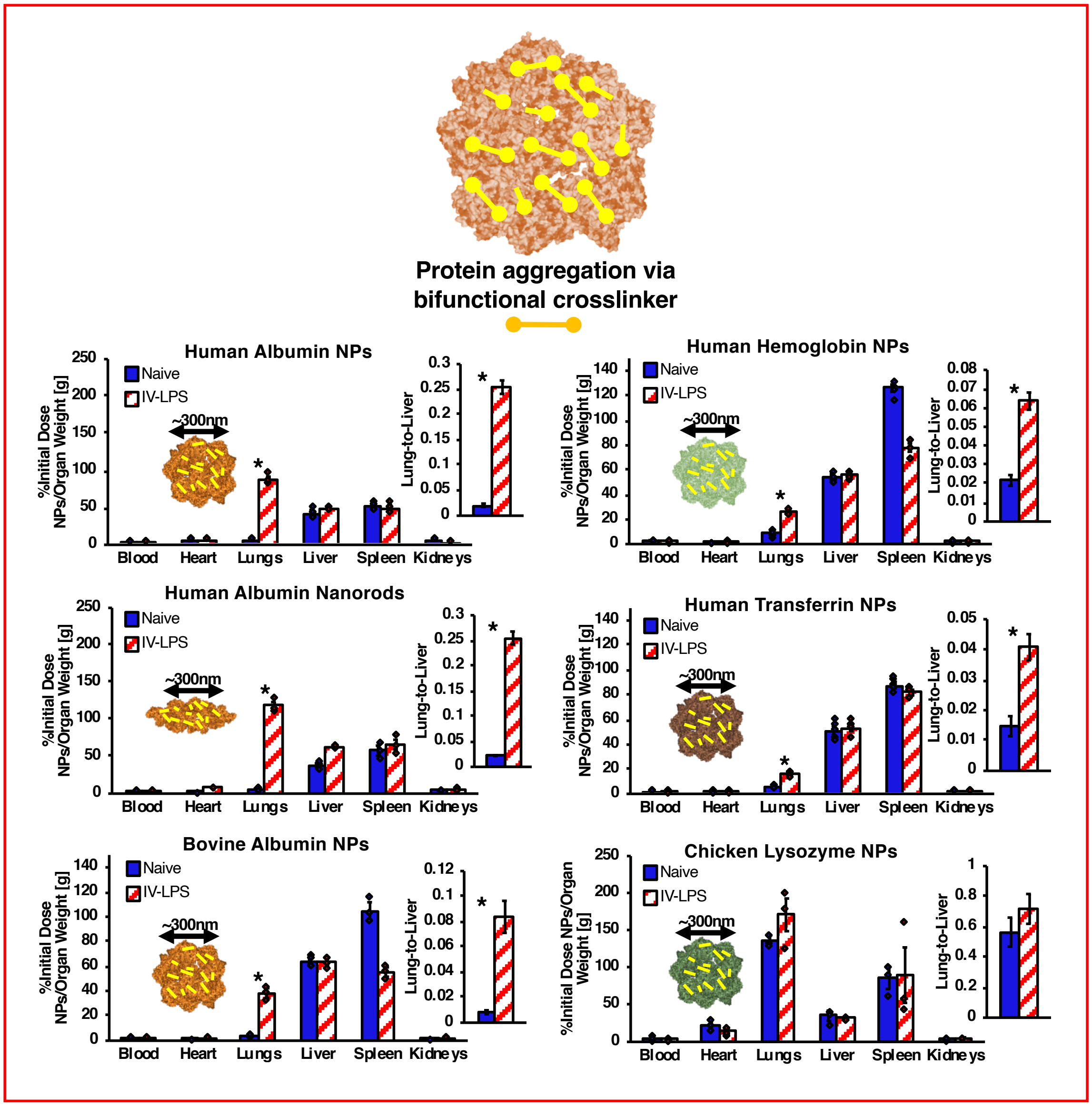


**Supplementary Figure 9**

*Biodistributions of Structural and Compositional Variants of Crosslinked Protein Nanoparticles in Naïve and IV-LPS-Injured Mice.* Different crosslinked protein nanoparticle formulations, as described in supplementary figure 1b, were traced in naïve and intravenous LPS-challenged mice. LPS treatment enhanced pulmonary nanoparticle uptake for all crosslinked protein nanoparticle variants, except for crosslinked lysozyme particles. For lysozyme particles, uptake in both injured and naïve lungs exceeded 20% of initial dose. Lung, liver, and blood data for human albumin nanoparticles, human albumin nanorods, and bovine albumin nanoparticles are reproduced in figure 3b in the main text. (** = p<.001*)

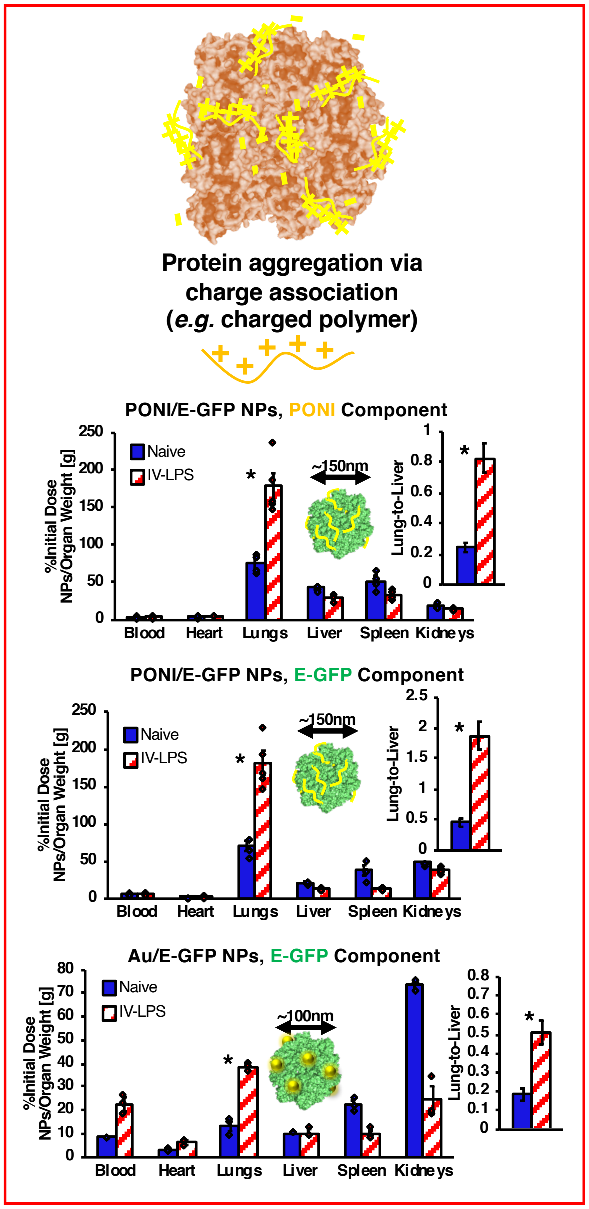


### Supplementary Figure 10

*Biodistributions of Compositional Variants of Charge-Agglutinated Green Fluorescent Protein Nanoparticles in Naïve and IV-LPS-Injured Mice.* Nanoparticles formed by combining glutamate-tagged green fluorescent protein with arginine-tagged gold nanoclusters (Au) or arginine-poly(oxanorborneneimide) (PONI) (see supplementary figure 1d) were traced in naïve and intravenous LPS-challenged mice. PONI-eGFP nanoparticles were traced by labeling eGFP with $^{125}$I and PONI with $^{131}$I. LPS treatment enhanced pulmonary nanoparticle uptake both types of particle. Simultaneous PONI and eGFP tracing indicated that both nanoparticle components localized to the lungs after LPS injury. Lung, liver, and blood data are reproduced in figure 3c in the main text. (* = *p<.001*)

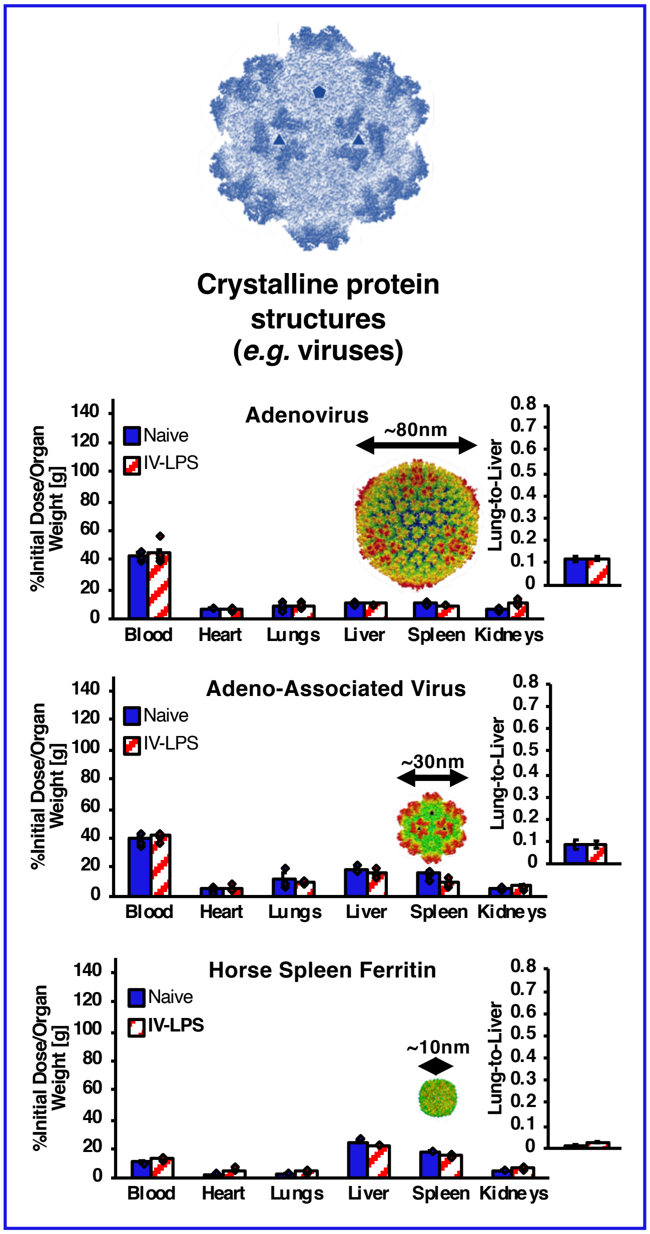


**Supplementary Figure 11**

*Biodistributions of Adenovirus, Adeno-Associated Virus, and Horse Spleen Ferritin Nanocages in Naïve and IV-LPS-Injured Mice.* Three naturally occurring crystalline protein nanostructures (see supplementary figure 1d for DLS data) were traced in naïve and intravenous LPS-challenged mice. LPS treatment had no effect on the biodistributions of radiolabeled adenovirus, adeno-associated virus and horse spleen ferritin. Lung, liver, and blood data are reproduced in figure 3d in the main text.

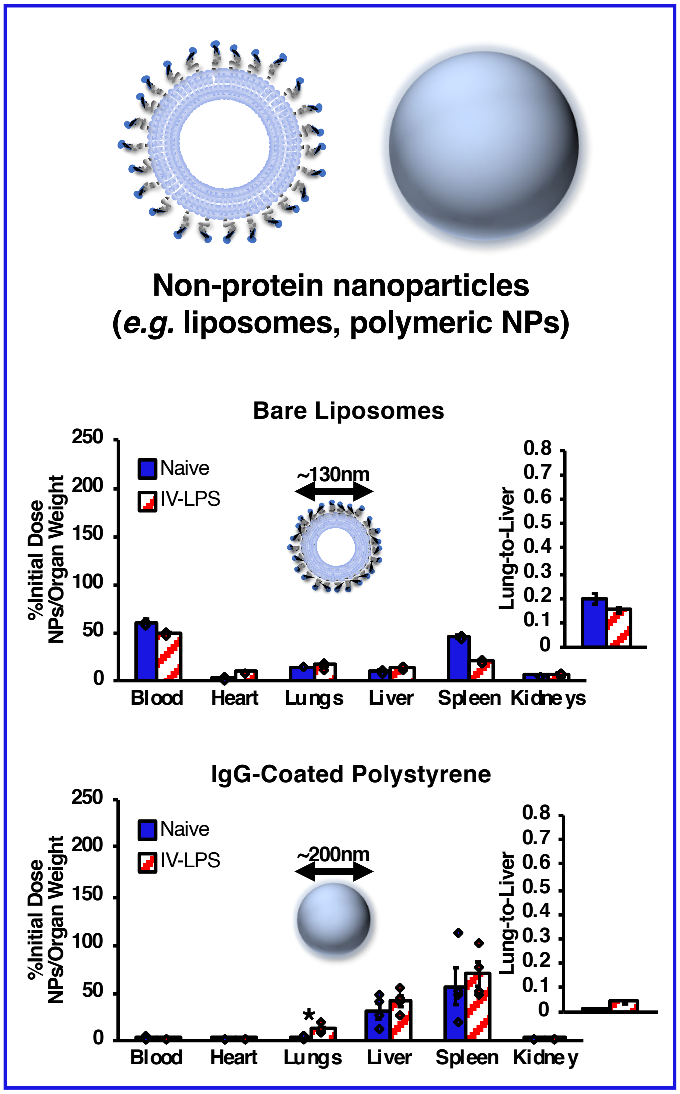


**Supplementary Figure 12**

*Biodistributions of Bare Liposomes and IgG-Coated Polystyrene Nanoparticles in Naïve and IV-LPS-Injured Mice.* As example nanoparticles not based on assembly of protein, bare liposomes (see supplementary figure 1c for DLS data) and IgG-coated polystyrene nanoparticles (see supplementary figure 1d for DLS data) were traced in naïve and intravenous LPS-challenged mice. LPS treatment had no effect on the biodistribution of bare liposomes. LPS treatment did enhance pulmonary uptake of IgG-coated polymeric nanoparticles, albeit with lower significance (** = p<.01*) and at lower levels of lung uptake than observed with variant nanogels, crosslinked protein particles, or charge associated protein particles. Lung, liver, and blood data are reproduced in figure 3e in the main text.

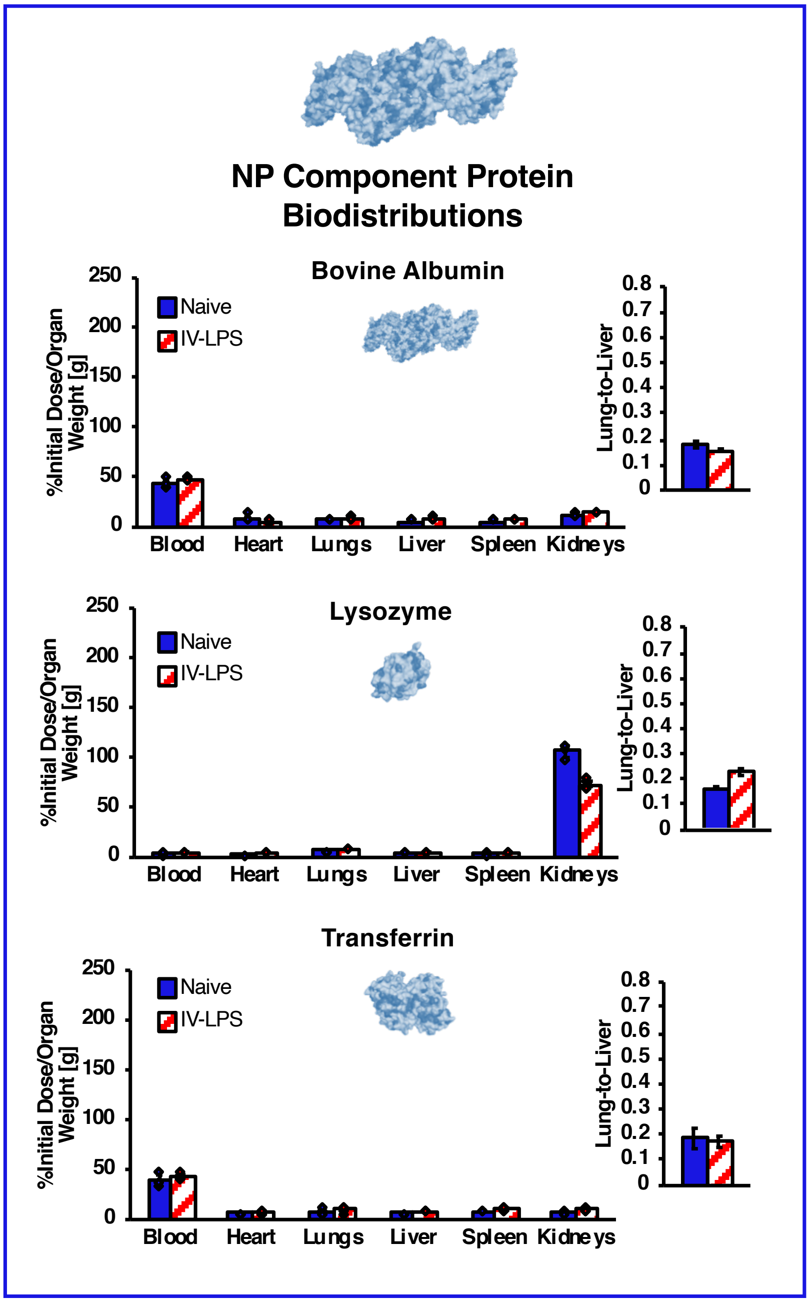


**Supplementary Figure 13**

*Biodistributions of Isolated Albumin, Lysozyme, and Transferrin in Naïve and IV-LPS-Injured Mice.* Different radiolabled isolated proteins were traced in naïve and intravenous LPS-challenged mice. LPS treatment had no effect on the biodistributions of bovine albumin, hen lysozyme, or human transferrin.

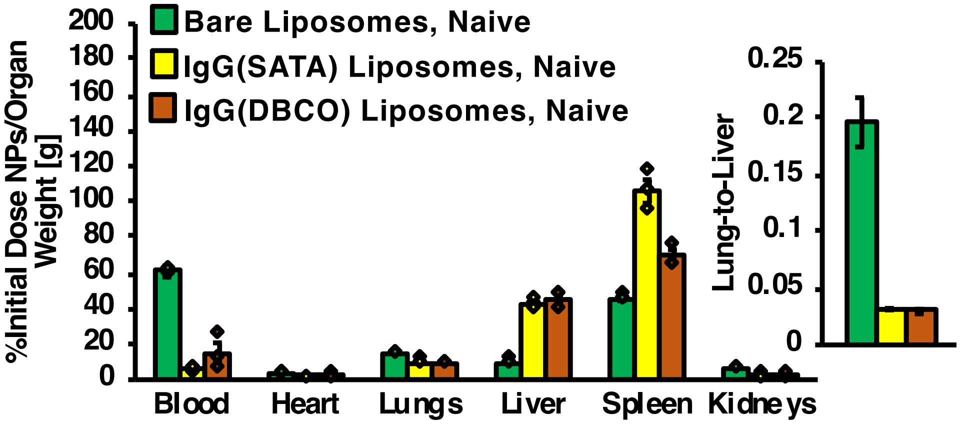


**Supplementary Figure 14**

*Biodistributions in Naïve Mice for Bare Liposomes, Liposomes Conjugated to IgG via SATA-Maleimide Reaction, and Liposomes Conjugated to IgG via DBCO-Azide Reaction.* In juxtaposition to biodistribution data in IV LPS-challenged mice, as presented in figure 4b in the main text, bare liposomes, IgG-SATA liposomes, and IgG-dibenzocyclooctyne (DBCO) liposomes were traced in naïve mice. Whereas IgG-DBCO liposomes had uniquely high levels of pulmonary uptake in LPS-challenged mice, no significant differences were noted in pulmonary uptake of the different liposome formulations in naïve mice.

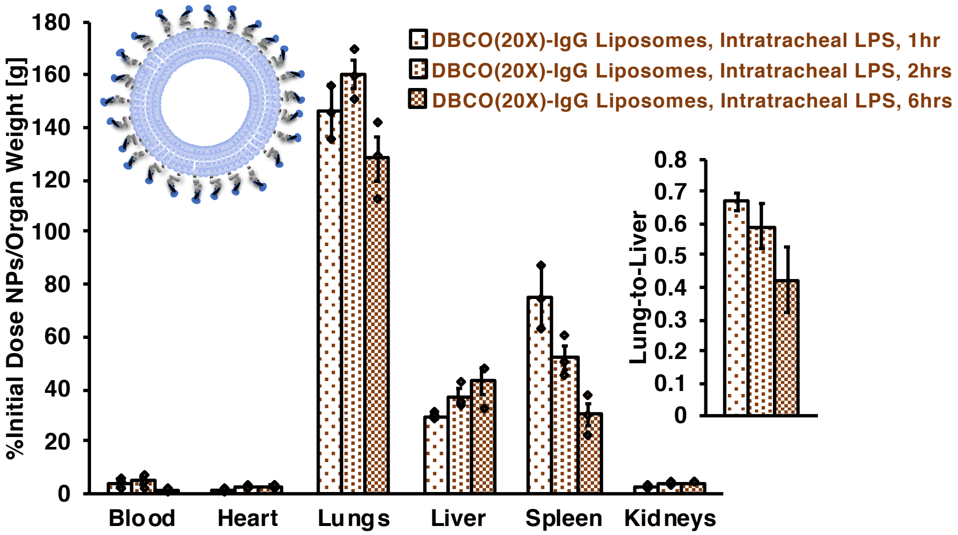


**Supplementary Figure 15**

*Biodistributions of DBCO:IgG (20:1) Liposomes in Mice 1, 2, and 6 Hours After Intratracheal LPS Injury.* As an alternative to intravenous LPS injection, mice were administered LPS via intratracheal (IT) instillation, prior to bolus dosing with liposomes coated with IgG conjugated to a 20-fold excess of DBCO. As with IV LPS-injured mice, IT LPS injury led to high levels of pulmonary uptake for DBCO(20X)-IgG liposomes. Similar levels of pulmonary uptake were observed at 1, 2, and 6 hours after IT LPS instillation, with liposomes circulating for 30 minutes for each data set.

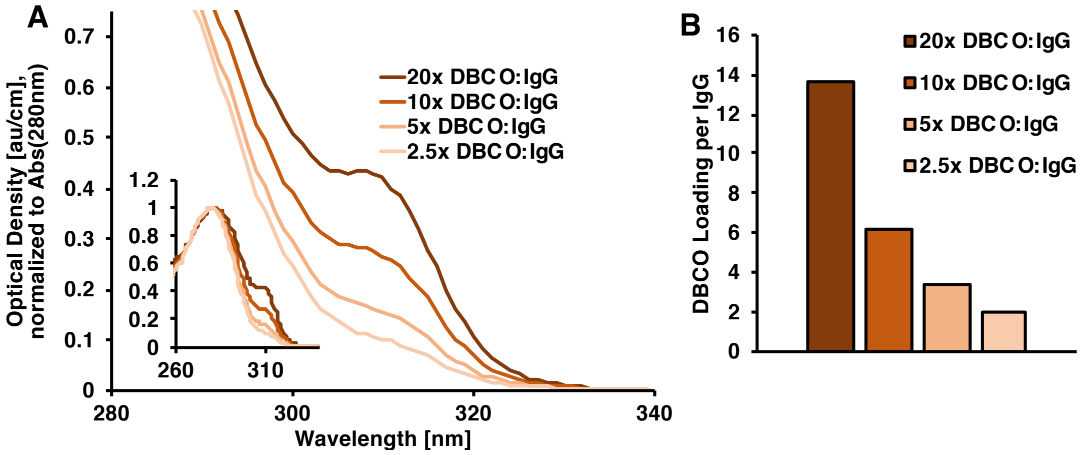


**Supplementary Figure 16**

*Spectrophotometric Characterization of DBCO Conjugation to IgG.* (a) IgG was reacted with 2.5-fold, 5-fold, 10-fold, and 20-fold excesses of DBCO-PEG$_4$-NHS ester and optical density of the resulting conjugates was determined between 220nm and 350nm. Absorbance at 309nm indicated DBCO on the IgG and absorbance at 280nm was indicative of IgG concentration (see inset for absorbance data at 280nm). (b) Spectral overlap of DBCO absorbance with IgG absorbance was noted by correcting absorbance at 280nm according to $Abs_{280C} = Abs_{280} - 1.089 \times Abs_{309}$. Molar IgG concentration was determined according to $[IgG] = \frac{Abs_{280C}}{\varepsilon_{280,IgG}}$, where $\varepsilon_{280,IgG}$ is the IgG extinction coefficient at 280nm, 204000 L mol$^{-1}$cm$^{-1}$. Molar DBCO concentration was determined according to $[DBCO] = \frac{Abs_{309}}{\varepsilon_{309,DBCO}}$, where $\varepsilon_{309,DBCO}$ is the DBCO extinction coefficient at 309nm, 12000 L mol$^{-1}$cm$^{-1}$. Number of DBCO per IgG was determined as the ratio $\frac{[DBCO]}{[IgG]}$.

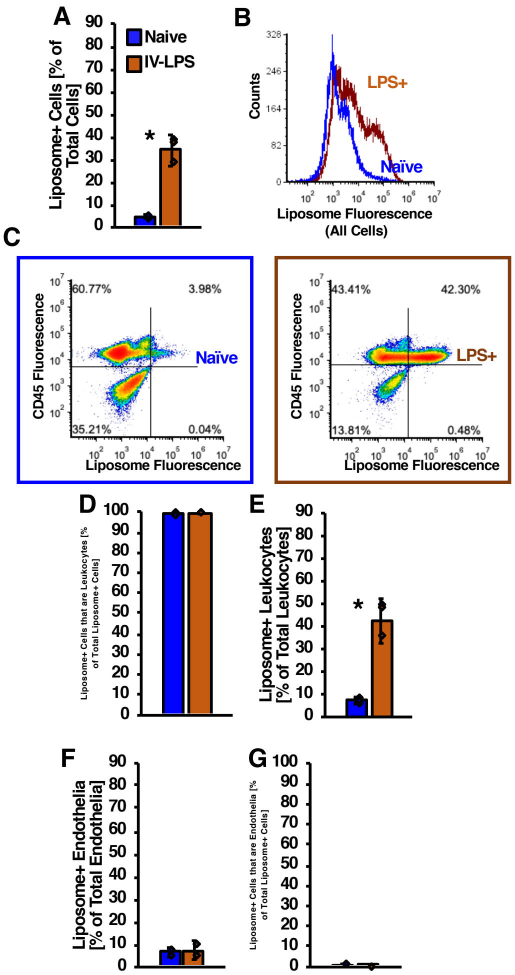


**Supplementary Figure 17**

*Flow Cytometric Characterization of DBCO:IgG (20:1) Liposome Uptake in Leukocytes and Endothelial Cells in Naïve and Inflamed Lungs.* (a) Fluorescence from DBCO(20X)-IgG liposomes containing green fluorescent TopFluor lipid was measured in single cell suspensions prepared from mouse lungs harvested after 30 minutes liposome circulation. With gates set as depicted in main text figure 4d, the number of cells positive for liposome fluorescence increased between naïve and LPS-challenged lungs. (b) A population of high-fluorescence cells was detected in IV LPS-challenged lungs, but not naïve lungs. (c) Fluorescence generated by CD45 staining, distinguishing leukocytes in single cell suspensions, plotted against DBCO(20X)-IgG liposome fluorescence in single cell suspensions prepared from naïve and IV LPS-challenged lungs. (d-e) With gates set by the quadrants delineated in (c), correlation between liposome fluorescence and CD45 staining indicated the percentage of liposome-bearing cells that were leukocytes as >95% in both naïve and IV LPS-challenged lungs. Similar analysis indicated that the fraction of leukocytes containing liposomes increased in LPS-challenged vs. naïve lungs. (f-g) Single cell suspensions were stained with CD31 antibody to indicate endothelial cells. Correlation between CD31 staining and liposome fluorescence indicated that <10% of endothelial cells contained liposomes and <1% of all liposome-positive cells in the suspensions were endothelial cells. (** = p<.01*)

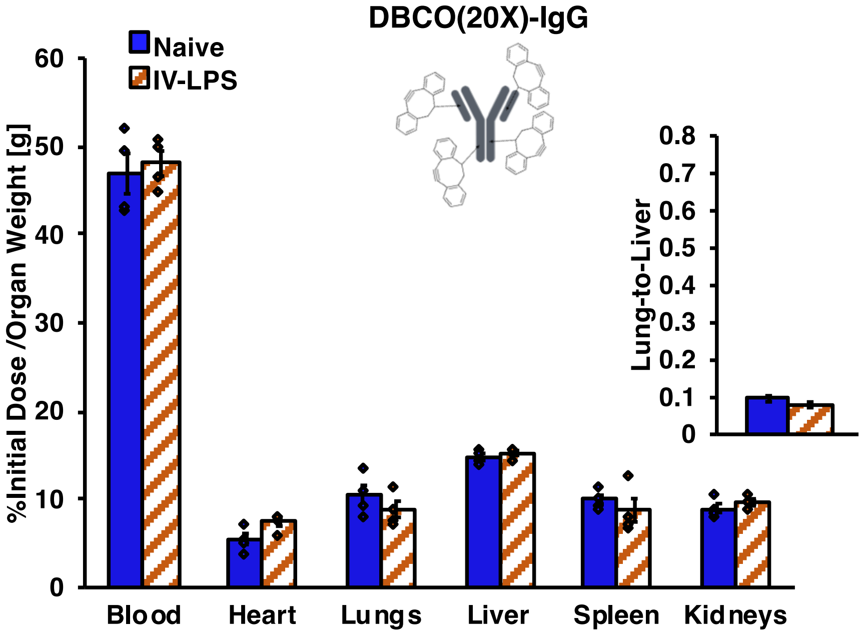


**Supplementary Figure 18**

*Biodistributions of Isolated DBCO:IgG (20:1) in Naïve and IV-LPS-Injured Mice.* IgG conjugated to a 20-fold excess of DBCO was traced in naïve and IV LPS-challenged mice. No significant differences were observed in isolated DBCO(20X)-IgG biodistributions between naïve and injured mice.

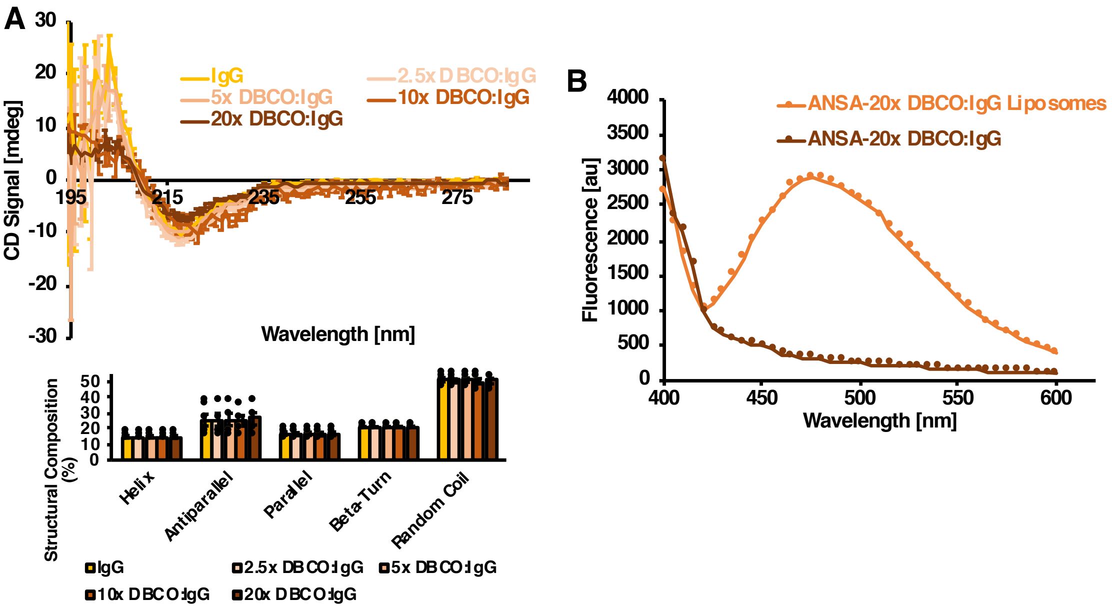


**Supplementary Figure 19**

*Circular Dichroism Spectroscopic Characterization of Protein Secondary Structure in DBCO-Modified IgG and ANSA Characterization of Hydrophobic Domain Accessibility on DBCO:IgG (20:1) Liposomes.* (a) Circular dichroism spectra for IgG modified with different concentrations of DBCO-PEG$_4$-NHS ester and unmodified IgG. Inset: neural network deconvolution of CD spectra indicating that IgG secondary structure composition was unchanged by modification with all tested densities of DBCO. (b) 8-anilino-1-naphthalenesulfonic acid (ANSA) staining of IgG modified with a 20-fold excess of DBCO or liposomes conjugated to DBCO(20X)-IgG, with free DBCO(20X)-IgG concentration matched to the DBCO(20X)-IgG concentration on the liposomes. Increased ANSA fluorescence indicates increased accessibility of hydrophobic domains on the liposomes, compared to free DBCO(20X)-IgG.

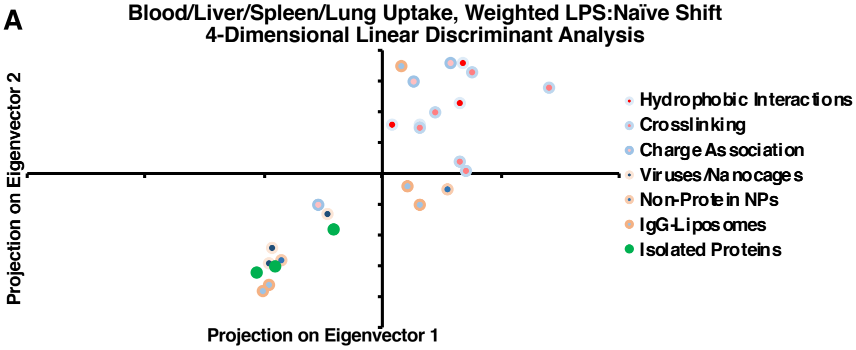


| | Eigenvector 1 | Eigenvector 2 | Eigenvector 3 | Eigenvector 2 |
|---|---|---|---|---|
| Blood | -0.28233 | -0.10194 | 0.59684 | 0.218224 |
| Liver | -0.32578 | 0.75672 | -0.25106 | 0.819177 |
| Spleen | -0.88632 | -0.18302 | 0.40374 | -0.528561 |
| Lungs | 0.16911 | 0.61927 | 0.64633 | 0.044156 |

| Eigenvalues |
|---|
| 3.881353 |
| 3.190054 |
| 0.225852 |
| 0.051049 |

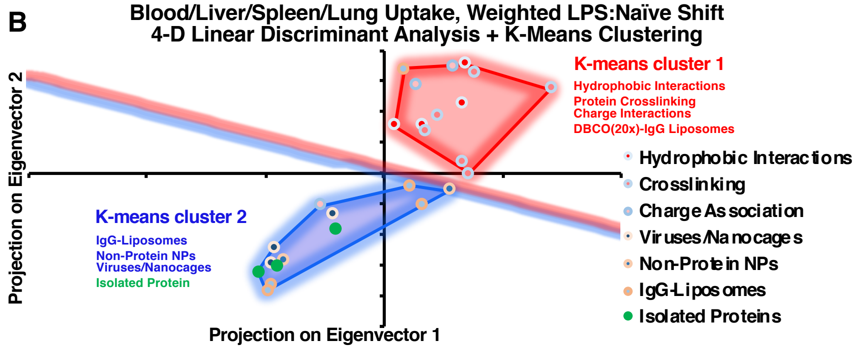


**Supplementary Figure 20**

*Linear Discriminant Analysis of Nanoparticle Biodistributions in Naïve and LPS-Injured Mice*. Nanoparticle and protein retention in blood, lungs, liver, and spleen in naïve and IV LPS-challenged mice was compiled for all tested nanoparticles and proteins. For each organ, weighted LPS:naïve shifts in nanoparticle uptake were determined according to $LPS{:}naive = (\overline{\%ID_{LPS}} - \overline{\%ID_{Naive}}) \times \frac{\overline{\%ID_{LPS}}}{\overline{\%ID_{Naive}}}$. LPS:naïve shift data for each organ were then centered and normalized via $LPS{:}naive_{norm} = (LPS{:}naive - \overline{LPS{:}naive})/\overline{LPS{:}naive}$. Centered and normalized LPS:naïve shifts were subjected to linear discriminant analysis, with data divided into subclasses defined as: hydrophobic interactions (nanogels), crosslinking (crosslinked protein nanoparticles), charge association (PONI-eGFP and Au-eGFP nanoparticles), viruses/nanocages (adenovirus, adeno-associated virus, and ferritin), non-protein NPs (polystyrene nanoparticles and bare liposomes), IgG-liposomes (encompassing SATA-maleimide and all variant DBCO conjugation chemistries), and isolated proteins. Projection of shift data along the first two eigenvectors is depicted in (a), with each eigenvector and corresponding eigenvalue enumerated in the inset table. Eigenvectors 1 and 2, accounting for >95% of variability in the data, were dominated by variation in pulmonary, hepatic, and splenic uptake. (b) Projections in (a) were subjected to K-means clustering analysis, indicating two clusters with lysozyme-dextran nanogels, DBCO(20X)-IgG liposomes, crosslinked protein nanoparticles, and glutamate-tagged GFP nanoparticles forming a single cluster.

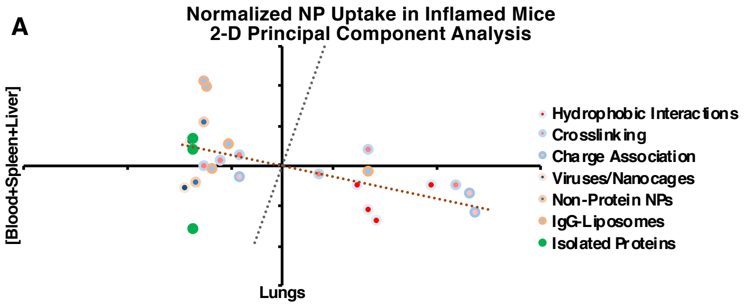


| | Eigenvector 1 | Eigenvector 2 | Eigenvalues |
|---|---|---|---|
| Lungs | -0.96438 | -0.26453 | 1.03398 |
| [Blood+Liver+Spleen] | 0.26453 | -0.96438 | 0.14179 |

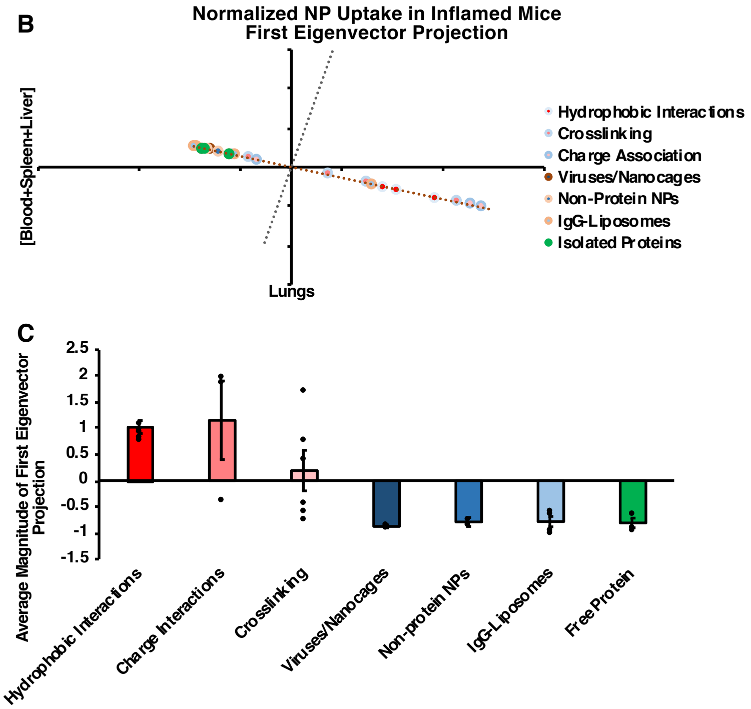


**Supplementary Figure 21**

*Principal Component Analysis of Nanoparticle Biodistribution Data in LPS-Injured Mice.* (a) Nanoparticle and protein retention in lungs vs. all other organs in IV LPS-challenged mice was compiled for all tested nanoparticles proteins. Data was centered and normalized according to $\%ID_{norm} = (\%ID - \overline{\%ID})/\overline{\%ID}$. Principal component analysis assessed eigenvectors depicted as dashed axes in (a) and enumerated in the inset table. (b) Centered and normalized data was projected along the first eigenvector. (c) Magnitude of the data projection along the first eigenvector was assessed and first eigenvector projection values were compiled in the nanoparticle classes described in supplementary figure 20, with DBCO(20X)-IgG liposomes excluded. Classes grouped into K-means cluster 1 (red/pink, see supplementary figure 20b) had significantly different first eigenvector projections, compared to classes grouped into K-means cluster 2 (blue/green).

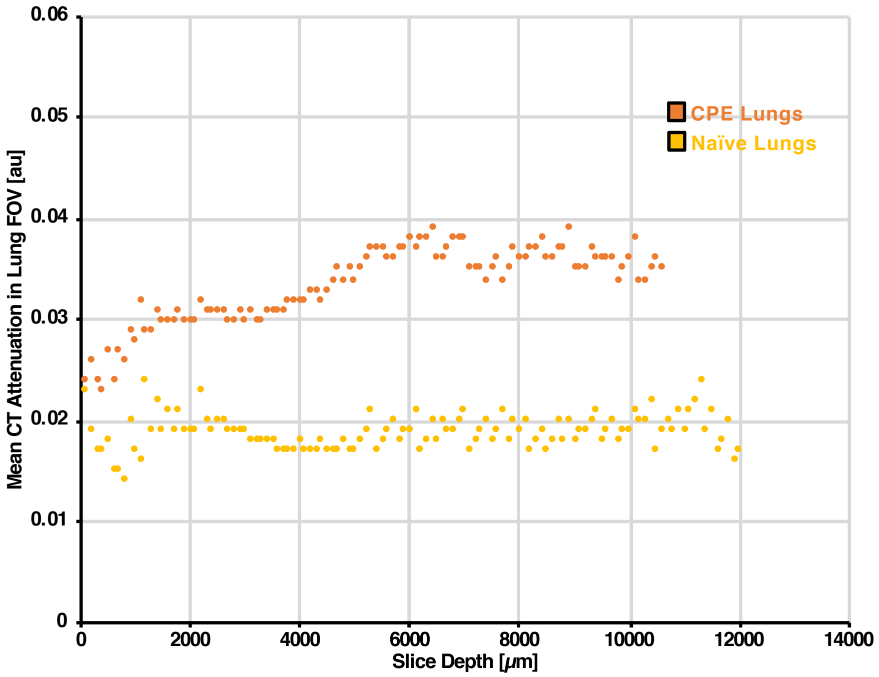


**Supplementary Figure 22**

*Quantification of CT Attenuation in Edematous and Naïve Mouse Lungs.* CT images, depicted in three-dimensional reconstructions in figure 5a in the main text, were obtained for a naïve mouse and a mouse afflicted with cardiogenic pulmonary edema. For each axial slice in the CT images, mean CT attenuation was determined in manually drawn fields of view encompassing the lungs. Mean CT attenuation is plotted above as a function of slice depth.

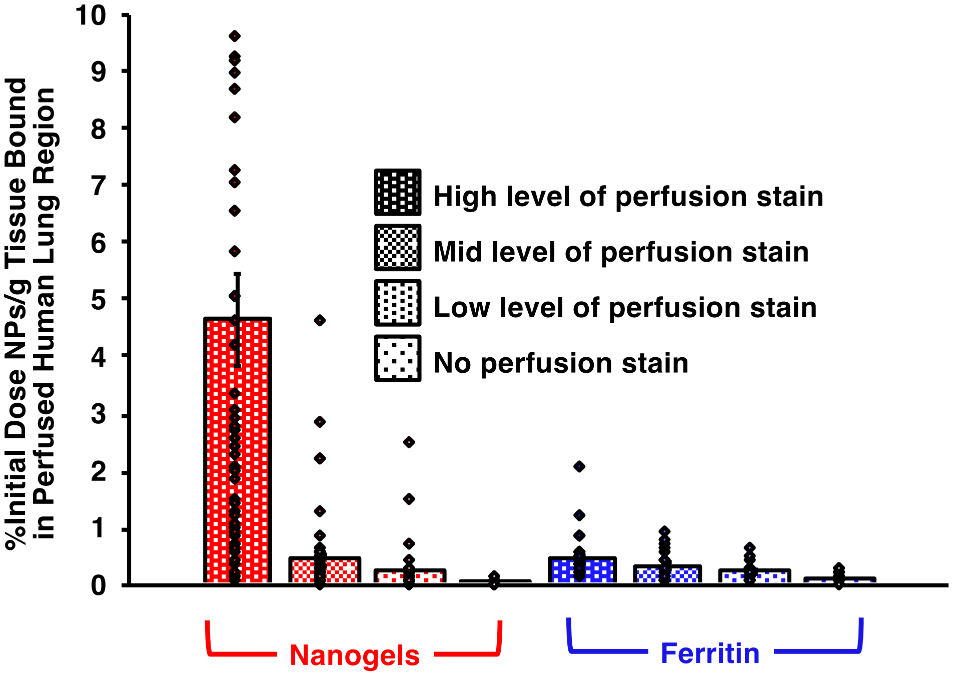


**Supplementary Figure 23**

*Lysozyme-Dextran Nanogel and Ferritin Nanocage Uptake in Human Lungs as a Function of Tissue Perfusion.* Human lungs were divided into ~1g segments according to levels of staining induced by tissue dye introduced via the same catheter used for nanoparticle administration, with staining divided by the experimenter into high, medium, and low levels of tissue staining. Nanoparticle retention in each segment was subsequently assessed by detection of radiolabel. Levels of lysozyme-dextran nanogel or ferritin retention in each type of segment are plotted above, indicating that nanogel retention is highly focused in directly perfused tissue

**Supplementary Table 1.** *Lysozyme-Dextran Nanogel and Crosslinked Albumin Nanoparticle Concentration in Lungs*

| Nanoparticle Concentration in Lungs | | |
|---|---|---|
| | Naïve | IV-LPS |
| Lysozyme-Dextran Nanogels | 5.25 %ID/g | 116.43 %ID/g |
| Albumin Nanoparticles | 6.34 %ID/g | 87.62 %ID/g |

**Supplementary Table 2.** *Flow Cytometric Assessment of Nanoparticle Specificity for Neutrophils*

| | Proportion of Neutrophils Containing Nanoparticles | | Proportion of Nanoparticle Uptake Accounted for by Neutrophils | |
|---|---|---|---|---|
| | Naïve | IV-LPS | Naive | IV-LPS |
| Lysozyme-Dextran Nanogels | 18.5% | 82.5% | 49.2% | 74.0% |
| Albumin Nanoparticles | 11.4% | 73.7% | 50.6% | 70.6% |

**Supplementary Table 3.** *Lysozyme-Dextran Nanogel Variants: Specificity for Injured Lungs*

| Lysozyme-Dextran Nanogels: Enhancement of Lung Uptake Following Injury | | | | |
|---|---|---|---|---|
| | 75nm LDNGs | 130nm LDNGs | 200nm LDNGs | 275nm LDNGs |
| Uptake in Injured Lungs | 122.27 %ID/g | 116.43 %ID/g | 156.05 %ID/g | 110.54 %ID/g |
| Lung Uptake (LPS):Lung Uptake (Naïve) | 16.32 | 22.18 | 12.82 | 4.94 |

**Supplementary Table 4.** *Crosslinked Protein Nanoparticle Variants: Specificity for Injured Lungs*

| Crosslinked Protein Nanoparticles: Enhancement of Lung Uptake Following Injury | | | | | |
|---|---|---|---|---|---|
| | Human Albumin NPs | Human Albumin Nanorods | Bovine Albumin NPs | Human Hemoglobin NPs | Human Transferrin NPs |
| Uptake in Injured Lungs | 87.62 %ID/g | 117.82 %ID/g | 39.42 %ID/g | 27.98 %ID/g | 16.73 %ID/g |
| Lung Uptake (LPS):Lung Uptake (Naïve) | 13.82 | 18.91 | 10.16 | 3.23 | 3.17 |

**Supplementary Table 5.** *Viruses and Nanocages Lack Specificity for Injured Lungs*

| Viruses and Nanocages: No Enhancement of Lung Uptake Following Injury | | | |
|---|---|---|---|
| | Horse Spleen Ferritin | Adeno-Associated Virus | 200nm LDNGs |
| Uptake in Injured Lungs | 4.45 %ID/g | 10.00 %ID/g | 8.94 %ID/g |
| Lung Uptake (LPS):Lung Uptake (Naïve) | 1.15 | 0.80 | 1.01 |

**Supplementary Table 6.** *Isolated Proteins Lack Specificity for Injured Lungs*

| Isolated Proteins: No Enhancement of Lung Uptake Following Injury | | | |
|---|---|---|---|
| | Bovine Serum Albumin | Hen Egg White Lysozyme | Human Transferrin |
| Uptake in Injured Lungs | 9.22 %ID/g | 8.92 %ID/g | 9.69 %ID/g |
| Lung Uptake (LPS):Lung Uptake (Naïve) | 1.19 | 1.31 | 1.10 |

**Supplementary Table 7.** *Cyclooctyne Copper-Free Click Chemistry Confers Immunoliposomes with Specificity for Injured Lungs*

| IgG Liposomes: Enhancement of Lung Uptake Following Injury for Copper-Free Click Liposomes | | | |
|---|---|---|---|
| | Bare Liposomes | SATA-IgG Liposomes | DBCO-IgG Liposomes |
| Uptake in Injured Lungs | 16.89 %ID/g | 22.26 %ID/g | 117.16 %ID/g |
| Lung Uptake (LPS):Lung Uptake (Naïve) | 1.14 | 1.63 | 11.89 |

**Supplementary Table 8.** *DBCO-IgG Liposome Specificity for Injured Lungs is Dependent on Cyclooctyne Concentration*

| Increasing DBCO:IgG = Increased Enhancement of DBCO-IgG Liposome Lung Uptake Following Injury | | | | |
|---|---|---|---|---|
| | DBCO(2.5X)-IgG Liposomes | DBCO(5X)-IgG Liposomes | DBCO(10X)-IgG Liposomes | DBCO(20X)-IgG Liposomes |
| Uptake in Injured Lungs | 16.91 %ID/g | 17.79 %ID/g | 31.35 %ID/g | 117.16 %ID/g |

**Supplementary Table 9.** *Flow Cytometric Assessment of DBCO-IgG Liposome Specificity for Neutrophils*

| Proportion of Neutrophils Containing DBCO-IgG Liposomes | | Proportion of Liposome Uptake Accounted for by Neutrophils | |
|---|---|---|---|
| Naïve | IV-LPS | Naive | IV-LPS |
| 9.7% | 49.5% | 48.4% | 88.5% |

**Supplementary Movie 1**
*Intravital Imaging of Lysozyme-Dextran Nanogel Uptake in IV-LPS-Injured Mouse Lungs.*

**Supplementary Movie 2**
*Three-Dimensional Reconstruction of Chest CT Images of Naïve Mouse Lungs.*

**Supplementary Movie 3**
*Three-Dimensional Reconstruction of Chest CT Images of Edematous Mouse Lungs.*

**Supplementary Movie 4**
*Three-Dimensional Reconstruction of SPECT-CT Imaging of Lysozyme-Dextran Nanogel Uptake in IV-LPS-Injured Mouse Lungs.*

**Supplementary Movie 5**
*Three-Dimensional Reconstruction of SPECT-CT Imaging of Lysozyme-Dextran Nanogel Uptake in Naïve Mouse Lungs.*

**Supplementary Movie 6**
*Three-Dimensional Reconstruction of SPECT-CT Imaging of Lysozyme-Dextran Nanogel Biodistribution in an IV-LPS-Injured Mouse.*

**Supplementary Movie 7**
*Three-Dimensional Reconstruction of SPECT-CT Imaging of Lysozyme-Dextran Nanogel Biodistribution in a Naïve Mouse.*

**4-Dimensional Linear Discriminant Analysis Gnu Octave Script**

```
%Input of normalized/weighted LPS:naïve shift data, definition of nanoparticle classes
XCnorm_all;
C=[1;1;1;1;1;2;2;2;2;2;2;3;3;3;4;4;4;5;5;6;6;6;7;7;7;7];

dimension=columns(XCnorm_all);
labels=unique(C);
C_LDA=length(labels);
Sw_LDA=zeros(dimension, dimension);
Sb_LDA=zeros(dimension, dimension);
mu_LDA=mean(XCnorm_all);
for i=1:C_LDA
   Xi_LDA=XCnorm_all(find(C == labels(i)),:);
   n_LDA=rows(Xi_LDA);
   mu_i_LDA=mean(Xi_LDA);
   XMi_LDA=bsxfun(@minus,Xi_LDA,mu_i_LDA);
   Sw_LDA=Sw_LDA + (XMi_LDA'*XMi_LDA);
   MiM_LDA=mu_i_LDA-mu_LDA;
   Sb_LDA = Sb_LDA + n_LDA*MiM_LDA'*MiM_LDA;
endfor
[W_LDA,D_LDA]=eig(Sw_LDA\Sb_LDA);
[D_LDA,i]=sort(diag(D_LDA),'descend');

%Output of eigenvectors and eigenvalues
W_LDA = W_LDA(:,i);
W_LDA(:,1)
W_LDA(:,2)
W_LDA(:,3)
W_LDA(:,4)
D_LDA

%Output of data projections along the different eigenvectors
XCnorm_all_proj_LDA=XCnorm_all*W_LDA(:,1:4)';
XCnorm_all_proj_LDA(:,1)
XCnorm_all_proj_LDA(:,2)
XCnorm_all_proj_LDA(:,3)
XCnorm_all_proj_LDA(:,4)
```

**2-Dimensional Principal Component Analysis Gnu Octave Script**

```
%Input of normalized LPS biodistribution data
XLnorm;

%Determine eigenvectors of the covariance matrix
CLnorm=cov(XLnorm);
[V,D]=eig(CLnorm);
[D,i]=sort(diag(D),'descend');

%Output of eigenvectors and eigenvalues
V=V(:,i)
D
cumsum(D)/sum(D)

%Output of data projections along the different eigenvectors
ZLnorm1=XLnorm*V(:,1);
ZLnorm2=XLnorm*V(:,2);
PLnorm1=ZLnorm1*V(:,1)';
PLnorm2=ZLnorm2*V(:,2)';
PLnorm1(:,1)
PLnorm1(:,2)
PLnorm2(:,1)
PLnorm2(:,2)
```
